# Dynamics and bifurcation structure of a mean-field model of adaptive exponential integrate-and-fire networks

**DOI:** 10.1101/2023.12.09.570909

**Authors:** Lionel Kusch, Damien Depannemaecker, Alain Destexhe, Viktor Jirsa

## Abstract

The study of brain activity spans diverse scales and levels of description, and requires the development of computational models alongside experimental investigations to explore integrations across scales. The high dimensionality of spiking networks presents challenges for understanding their dynamics. To tackle this, a mean-field formulation offers a potential approach for dimensionality reduction while retaining essential elements. Here, we focus on a previously developed mean-field model of Adaptive Exponential (AdEx) networks, utilized in various research works. We provide a systematic investigation of its properties and bifurcation structure, which was not available for this model. We show that this provides a comprehensive description and characterization of the model to assist future users in interpreting their results. The methodology includes model construction, stability analysis, and numerical simulations. Finally, we offer an overview of dynamical properties and methods to characterize the mean-field model, which should be useful for for other models.

## 1 Introduction

The study of brain activity is a complex and multifaceted field that involves examining different scales and levels of description. In addition to experimental work in neuroscience, computational models have been developed to investigate brain activity at different scales, ranging from molecular to whole-brain levels. These models of different natures can have different functions, such as explicative or predictive. In this paper, we are interested in models reproducing dynamical aspects of neuron membrane excitability, networked through a synaptic communication model. These networks are made up of an excitatory population and an inhibitory population in similar proportions to what is observed in the cortex. These networks are useful for studying complex dynamics associated with the large dimensions of these systems [Carlu et al., 2020, Depannemaecker et al., 2022]. However, having a large number of dimensions can be a limiting factor for the understanding of the representation of its dynamics [Depannemaecker et al., 2023]. But also, when going to a larger spatial scale to constitute a network model of cerebral regions [Goldman et al., 2020], if keeping this level of description, the system becomes too complex to be understood and too computationally demanding to be simulated. In these cases, it may be interesting to try to reduce the dimensions of the system, trying to keep key elements to achieve the function of the model. A possible way of reduction is the formulation of a mean-field. The present study focuses on the study of a mean-field previously developed [di Volo et al., 2019a, El Boustani and Destexhe, 2009, Zerlaut and Destexhe, 2017] and already used in different studies [Capone et al., 2019, Carlu et al., 2020, Chemla et al., 2019, di Volo et al., 2019b, Di Volo and Férézou, 2021, Goldman et al., 2023, Goldman et al., 2020, Tesler et al., 2022, Tesler et al., 2023, Zerlaut et al., 2018]. Such approaches may focus on the proper dynamics of its corresponding spiking network [Capone et al., 2019, Carlu et al., 2020, di Volo et al., 2019b, El Boustani and Destexhe, 2009] or on experimental results [Chemla et al., 2019, Di Volo and Férézou, 2021, Zerlaut et al., 2018] or, to clinically observable measures [Goldman et al., 2023,Tesler et al., 2022,Tesler et al., 2023]. However, despite a wide range of uses, no systematic study of the mean-field model properties was published previously. Therefore, we propose a description and characterization intended for future users of this model and thus to help them interpret their results. We propose a methodology, and the results are given for one specific mean-field derivation (i.e. on a specific fit of the transfer function, see models and methods section); each user may need to apply these approaches to characterize its specific model. In the “model and methods” section, we detail the construction of this model, and then in the “stability analysis” section, we detail important elements of the structure underlying the dynamics. Then, we propose a characterization of its implementation in numerical simulations. Finally, in a discussion, we situate this model in the broad panorama of mean-fields, explaining its advantages and limits.

## 2 Models and methods

In this section, we first describe the Adaptive Exponential Integrate-and-Fire model [Brette and Gerstner, 2005] that constituted the spiking network, and then the corresponding mean-field model. The equations’ details and the parameters’ values are available in Supplementary Tables 1 and 2 based on the proposition of Nordlie et al. [Nordlie et al., 2009].

### 2.1 AdEx network

A network of 10 000 spiking neurons is constructed with a probability of connection of 5%, according to a sparse and random (Erdos-Renyi type) architecture. The network consists of 20% inhibitory (FS) population, and 80% an excitatory (RS) population, accordingly to the proportions in cerebral cortex, as previously modelled [Carlu et al., 2020, Zerlaut et al., 2018]. The dynamics of a single neuron membrane excitability is described by the AdEx model [Brette and Gerstner, 2005] corresponding to the equation (1)

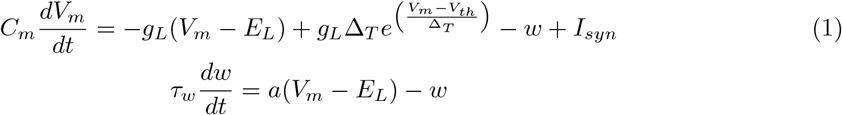

When an action potential is emitted (i.e. the membrane potential crosses a threshold), the system is reset as in the equation (2):

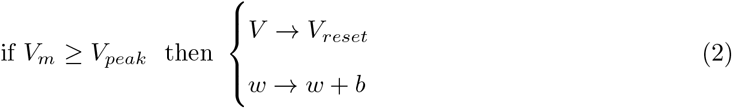

The interaction between AdEx neurons occurs through synaptic conductances, given by:

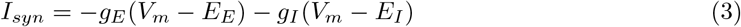

Where *E*_*E*_ = 0 mV and *E*_*I*_ = −80 mV are, respectively, the reversal potentials of excitatory synapses and inhibitory synapses. For each incoming spike, the excitatory and inhibitory conductances *g*_*E*_ and *g*_*I*_ are increased respectively by the value *Q*_*E*_ = 1.5 nS and *Q*_*I*_ = 5 nS. The conductances decrease following an exponential decay with a time constant *τ*_*syn*_ = 5 ms according to the equation (4).

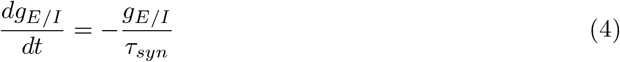

The network is simulated using the simulator NEST [Hahne et al., 2021].

### 2.2 AdEx Mean-field

Following the formalism proposed previously [El Boustani and Destexhe, 2009], di Volo and colleagues derived a mean-field model of AdEx networks with adaptation [di Volo et al., 2019a]. The differential equations describe the time evolution of the firing rate *ν*_*μ*_ of each population *μ* = *e, i* (equation (5)), the covariance *c*_*λη*_ between population *λη* (equation (6)), and, of the average adaptation for the excitatory population *W* (equation (7)).

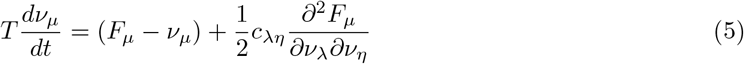

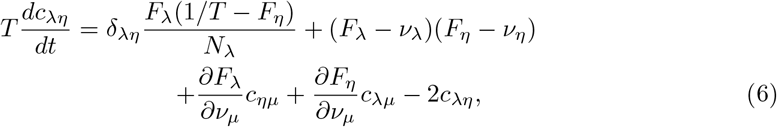

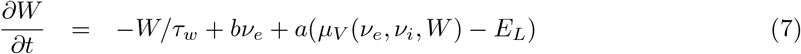

The parameter *T*, the time constant of the firing rate equations and covariance equations, comes from an essential assumption for this derivation: the network dynamics is considered as Markovian within a given time resolution T [El Boustani and Destexhe, 2009]. It is thus considered that each neuron emits a maximum of one spike into the time step T. Therefore, the value of T strongly constrains the maximum firing rate that can be considered according to these hypotheses. This question will be addressed in the discussion.

The parameters *b, a* et *E*_*L*_ in the equation (7), directly correspond to the parameters named the same in the AdEx model [di Volo et al., 2019a] ((1)).

The originality of this mean-field approach is that it is specified by the transfer functions *F*_*e,i*_, which expresses the firing rate of the population as a function of its inputs. These functions can be estimated using a semi-analytical approach [Zerlaut et al., 2018], consisting of fitting single-cell simulations with an analytic template. The transfer functions *F*_*e,i*_ for excitatory and inhibitory populations generally depend on three sources of input, *F*_*e*_(*ν*_*e*_ + *ν*_*ext*_, *ν*_*i*_), where *ν*_*ext*_ corresponds to a constant value of the firing rate an external drive. To evaluate the transfer functions, the first step is to measure the output firing rate of the single neuron model while receiving excitatory and inhibitory inputs. These measures are necessary because, due to the nonlinearity of the dynamics of the single neuron model, it cannot be inferred analytically, as well as determining when the system will irretrievably produce a spike. Thus, it is considered an effective or phenomenological threshold, 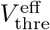, a function of the statistics of the subthreshold membrane voltage dynamics. This statistic is assumed to be normally distributed, with the average membrane voltage *μ*_*V*_, its standard deviation *σ*_*V*_ and auto-correlation time *τ*_*V*_ ; their calculation is described below. The effective threshold function is considered as a second-order polynomial giving the equation (8), where 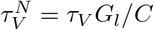. The {*P*} values are computationally found through fitting methods.

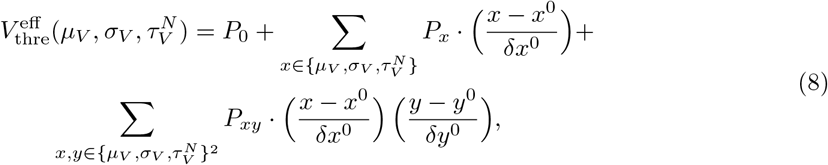

According to the semi-analytic approach, we fit the transfer function (equation (9)), where *erfc* is the Gauss error function, and thus obtain the output firing rate.

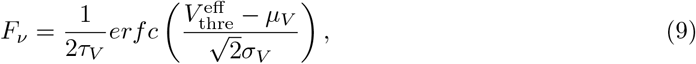

Considering asynchronous irregular regimes, we assume that the input spike trains follow Poissonian statistics [di Volo et al., 2019a,El Boustani and Destexhe, 2009,Zerlaut and Destexhe, 2017] and we calculate the averages (*μ*_*Ge,Gi*_) and standard deviations (*σ*_*Ge,Gi*_) of the conductances in the equations 10. In these equations, *K*_*e*_ and *K*_*i*_ are the average input connectivity received respectively from the excitatory or inhibitory population. As in the spiking network, *τ*_*e*_ = *τ*_*i*_ = *τ*_*syn*_ are synaptic time constants, and *Q*_*e*_ and *Q*_*i*_ the quantal increment of the conductances, respectively, for the excitatory or inhibitory populations.

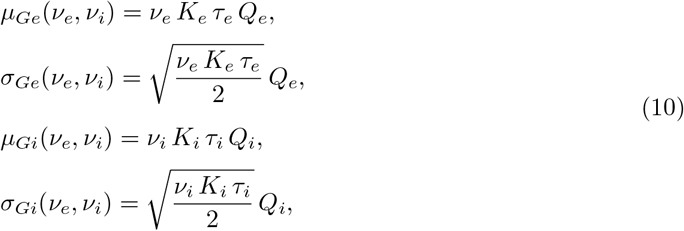

We can then calculate the total input conductance of the neuron *μ*_*G*_ and its effective membrane time constant 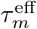 (equations (11)), with *C* and *g*_*L*_ being the same as in the AdEx model.

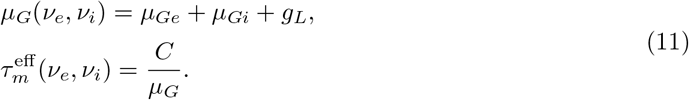

Then, we can calculate, in the equation (12), the mean subthreshold voltage assuming that the subthreshold moments (*μ*_*V*_, *σ*_*V*_, *τ*_*V*_) are not affected by the exponential term of the AdEx model, and thus considering only the leakage term and the synaptic inputs.

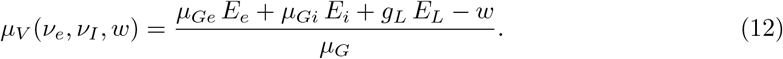

Thanks to the calculation developed by Zerlaut and al. [Zerlaut et al., 2018], we obtain *σ*_*V*_ and *τ*_*V*_ in the equations (13) and (14), where 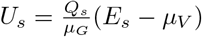 and *s* = (*e, i*).

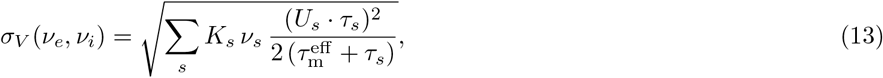

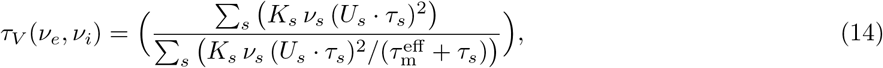

The mean-field is simulated using The Virtual Brain [Sanz-Leon et al., 2015].

### 2.3 Transfer function fitting

The transfer function is a function that provides the firing rate of a population for a given excitatory and inhibitory input firing rates and mean adaptation current. To fit the phenomenological threshold of this function, we create a data set of averages of the mean firing rate of 10 seconds of fifty neurons connecting only to one excitatory and one inhibitory Poisson generator with a fixed negative input. We take twenty values for inhibitory input firing rate (between 0 and 40 Hz) and twenty values of adaptation current, i.e. a constant negative current (between 0 and 500 pA). The five hundred excitatory input firing rates (between 0. and 200 Hz) are adjusted to get a precision of the output firing rate (the interval between two values is in the range of 0.1 and 1.0 Hz). The mean output firing rate is the average over fifty neurons by removing the outliers (values higher than three times the interquartile difference). The phenomenological voltage threshold of the transfer function is fitted following the paper of Zerlaut et al. 2018 [Zerlaut et al., 2018] with two steps. The initial values of the ten polynomial coefficients of the voltage threshold are set to (effective mean voltage, 1e-3, 1e-3, 1e-3, 1e-3, 0, 0, 0, 0, 0). The first fitting step is the minimization of mean square difference of the voltage threshold estimated from the transfer function 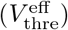 and the estimated effective threshold from the data set 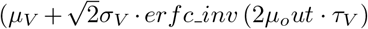). The second fitting step is the minimization of the mean squared difference between the output firing rate of the data set and the one estimated by the transfer function. The coefficients fit using Broyden–Fletcher–Goldfarb–Shanno algorithm [BROYDEN, 1970, Fletcher, 1970, Goldfarb, 1970, Shanno, 1970] implemented in Scipy [Virtanen et al., 2020].

### 2.4 Bifurcation analysis

The bifurcation analysis of the mean Ad Ex has been done using MatCont (version 7.7) [Dhooge et al., 2003], a numerical continuation library of MATLAB [Inc., 2022]. From the equilibrium point of a silence network (0.0, 0.0, 0.0, 0.0, 0.0, 0.0), we compute the change of stability depending on the external excitatory firing rate. The result of the numerical continuation is a stability diagram of the mean-field with a bifurcation point. The steps of numerical continuation are variable and adjusted depending on the sensitivity of the stability of the fixed-points.

### 2.5 Spike train analysis

In this paper, we analyze the spike trains analysis to validate the mean-field hypotheses and to compare the response to oscillatory input between the network and the mean-field. The hypotheses of the mean-field are evaluated using two measures: the time scale of the spike trains (auto-correlation time) and the minimum of the spiking interval.

The auto-correlation time is calculated based on the following equation (15) proposed by Wieland et al. 2015 [Wieland et al., 2015]:

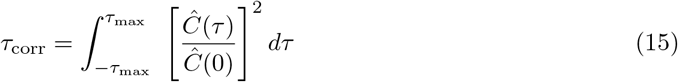

where *Ĉ*(*τ*) = ⟨*x*(*t*)*x*(*t* − *τ*)⟩ ⟨*x*⟩^2^ − ⟨*x*⟩ *δ*(*τ*) with 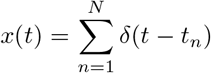 for *t*_*n*_ is the n-th spike time of the spike trains.

The minimum inter-spiking interval is the minimum time between two spikes of the same neurons. The maximum firing rate is the maximum number of spikes in 5 milliseconds.

The analysis of the network’s response to a sinusoidal Poisson generator is based on quantifying the phase difference between the oscillation of the mean firing rate of the excitatory neuron of the spiking network and the input oscillation. The network oscillation signal corresponds to the firing rate filter at the input oscillation frequency. The firing rate is defined as a sliding average with a window of 5 milliseconds of the instantaneous firing rate (resolution 0.1ms).

The phase difference between these two oscillatory signals is defined by the following equation(16):

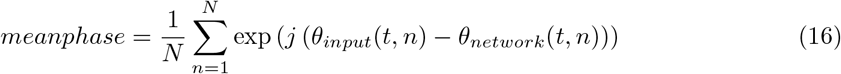

where *θ*_*input*_ is the analytic signal given by the Hilbert transformation of the sinusoidal input, and *θ*_*network*_ is the analytic signal provided by the Hilbert transformation of the network oscillation signal. The network oscillation signal is the histogram of instantaneous firing rate (bin:0.1) smooths with a sliding average window of 5 milliseconds. The absolute value of the *meanphase* corresponds to the phase locking value as defined by Lachaux et al. 1999 [Lachaux et al., 1999]. The angle of the *meanphase* corresponds to the phase shift [Freeman et al., 2003]. Additional analyses are described in the Supplementary Note 3.

### 2.6 Mean-field analysis

The mean-field signal corresponds to the excitatory mean firing rate filter at the input oscillation frequency. The rest of the analysis is the same as for the spike trains. The difference of phase is calculated based on the equation (16) where *θ*_*input*_ is the analytic signal given by the Hilbert transformation of the sinusoidal input and *θ*_*mean*_*field*_ is the analytic signal given by the Hilbert transformation of the mean-field signal. The phase locking value and the phase shift are extracted from this phase and used to quantify the mean-field’s response. Additionally, the maximum firing rate of the mean-field corresponds to this maximum value of the mean firing rate. Additional analyses are described in the Supplementary Note 4.

## 3 Results

In the results sections of this paper, we do not present specific simulations but instead, provide a characterization of the model itself. The results presented (unless otherwise indicated) are based on a unique set of parameters that have led to a fixed transfer function fitted for each of the populations (inhibitory and excitatory) considered. These results may differ for other fits and parameters, and we invite users of the mean-field AdEx to perform their own characterizations for their specific applications.

We start by showing the steady-state solutions of the AdEx mean-field.

### 3.1 Steady-state and transfer function

By construction, the AdEx mean-field (MF) model captures the out-of-equilibrium asynchronous irregular steady-state of the spiking neural network as a stable fixed-point. The stability analysis, according to the parameter *ν*_*ext*_, the external input, provides a bifurcation diagram. This bifurcation diagram can complete the understanding of the underlying structure of attractors while building a network of MF [Goldman et al., 2020, Zerlaut et al., 2018].

A qualitative comparison of the mean firing rate for a subset of external input characterizes the link between the bifurcation diagram of the MF and the spiking neural network state. The external input in this paper is a parameter, constant input, for the MF and the frequency of the Poissonian spike generator for the network. Figure 1 illustrates in the central plot the stability of the MF (black line) for different levels of external inputs and the corresponding mean firing rate of the steady-state of the network. The bifurcation diagram shows three regions of stability associated with the two variables, the mean firing rate of each population, also existing for the network. The side figures of the diagram show two simulations corresponding to two examples taken from the stable region of the MF with the associated network simulations. The qualitative comparison of these two measures displays a similar behavior and proves that the MF captures well the steady state of the network for different inputs. Remarkably, in the region with a very high firing rate (≈ 200Hz, the network is no longer in an asynchronous irregular regime, but the mean firing rate is captured by the MF. Furthermore, the qualitative comparison for different values of the spike-triggered adaptation indicates that the mean field captures fit the steady-state of the network in the presence of the adaptation current (see figure 2). However, looking more into details, significant quantitative differences appear for certain values of external inputs.

**Figure 1:**
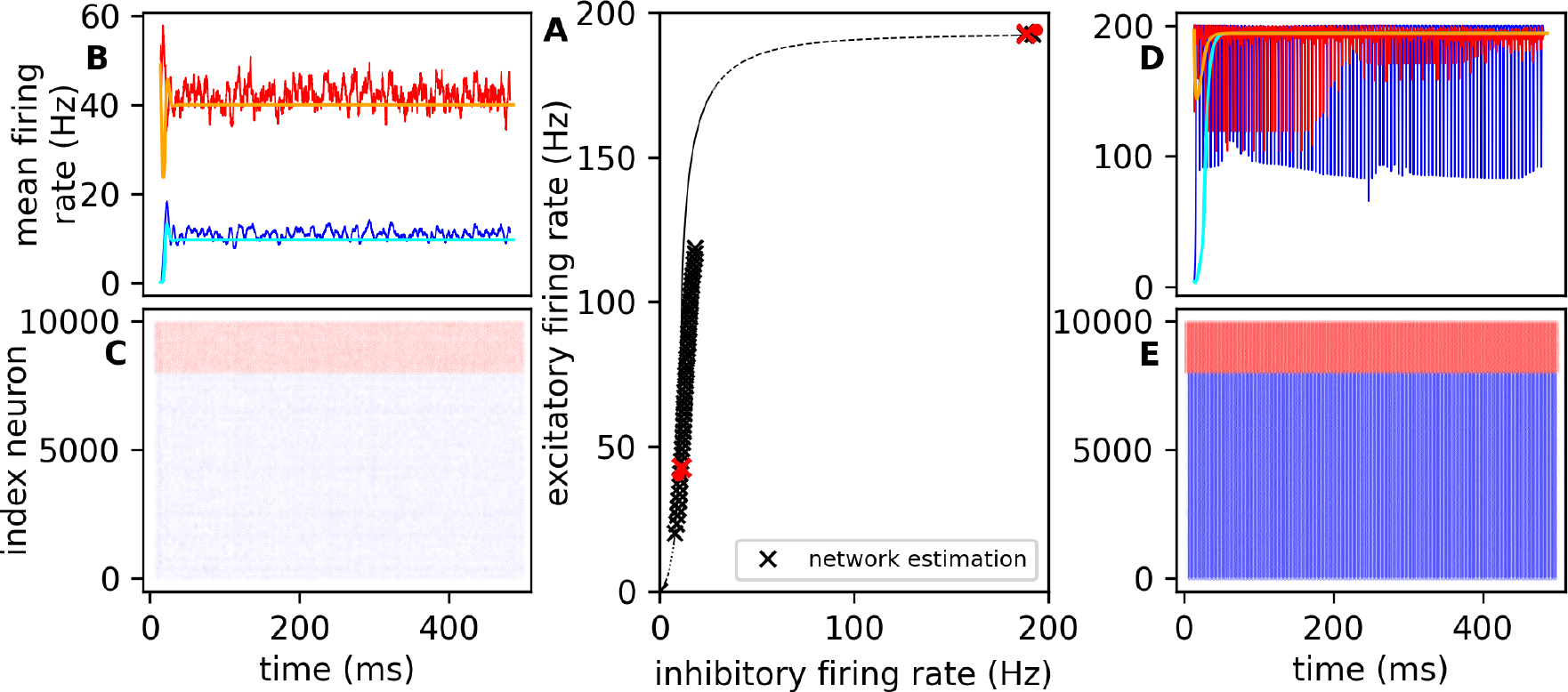
Mean-field and network dynamics without adaptation. Panel **A** represents the bifurcation of external excitatory input in the variable space of excitatory and inhibitory mean firing rate. The black line is the mean-field bifurcation with its stability (continuous line for stable fix points and dashed lines for unstable fix points). The crosses are the estimation of the fixed-point using the simulation of the spiking neural network. The red crosses and red points are the fixed-points associated with the examples of each side. Panels **B**,**C**,**D** and **E** exemplify network and mean-field simulations with an external excitatory input of 10 Hz and 80 Hz. The top panels (B, D) visualize the mean firing rate of the excitatory (red) and inhibitory (blue) neurons from the network simulation and the mean firing rate of the excitatory (orange) and inhibitory (cyan) population from the mean-field simulation. The bottom panel (C, E) are the corresponding raster plots of the spiking network simulations. The figures B and C display asynchronous irregular regime, and the figures D and E display synchronous regular regime

**Figure 2:**
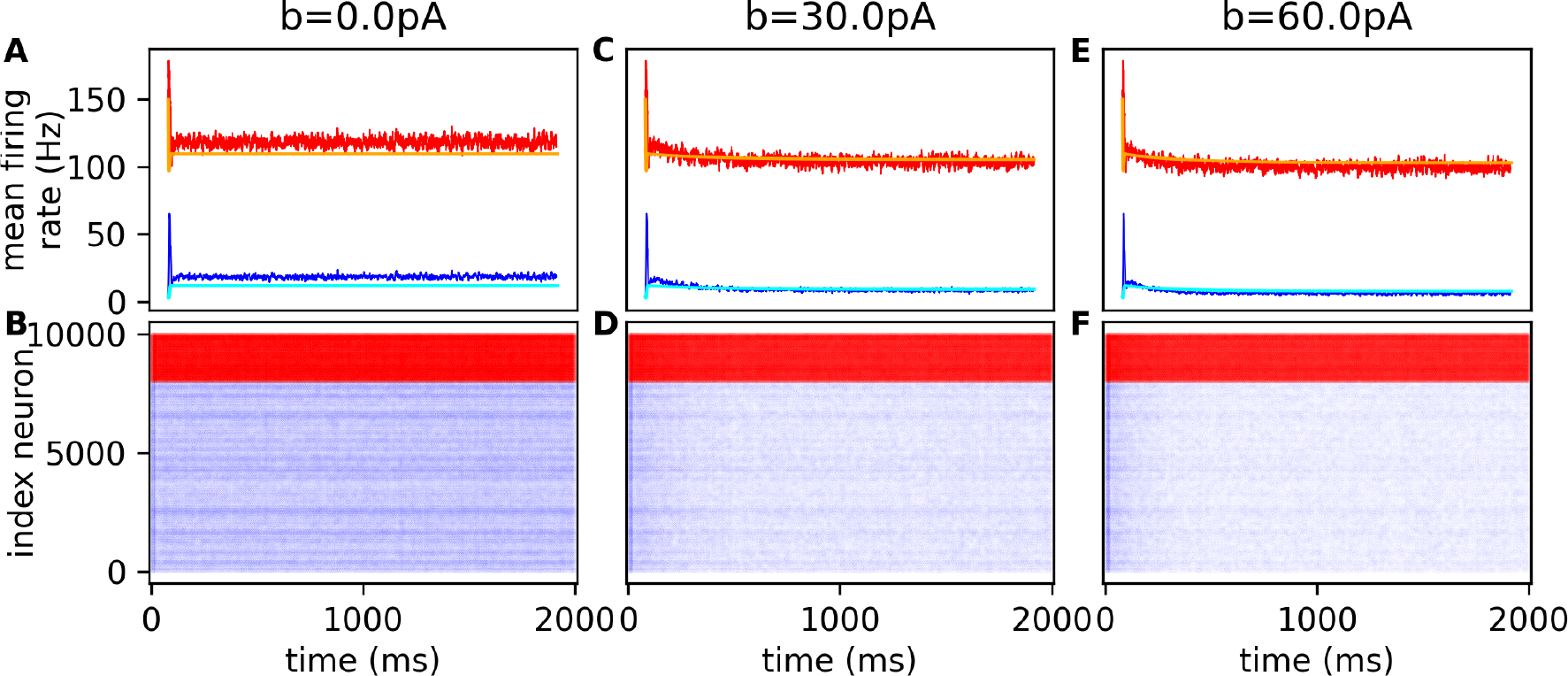
Mean-field and network dynamics with adaptation. Panels **A, C**, and **E** compare the mean firing rate of the network and mean-field simulation with an external excitatory input of 50Hz for different values of spike-triggered adaptation, respectively 0.0 pA, 30.0 pA and 60.0 pA. Panels **B, C**, and **F** are the corresponding raster plots of the spiking networks simulation.

Before looking into details, it is important to notice that the measure of the mean and standard deviation of the firing rate of a network’s steady-state present variability in time and is dependent on the window size. The supplementary figures 5 and 6 show the distribution of these measures over thirty seconds for different window sizes (for details see supplementary note 1). This confirms that the distribution is dependent on the network state. To reduce the measure variability, the steady-state of the network is measured over four seconds after discarding one second of transient. Figure 3 compares MF fixed-points and the corresponding mean firing rate of steady-state of the network. The results show the firing rate of each population for three different values of the parameter *b*, which directly affects the range of the adaptation variable *w*. This adaptation variable is a slower variable playing an important role in modeling phenomena such as neuromodulation [Goldman et al., 2020, Tort-Colet et al., 2023]

**Figure 3:**
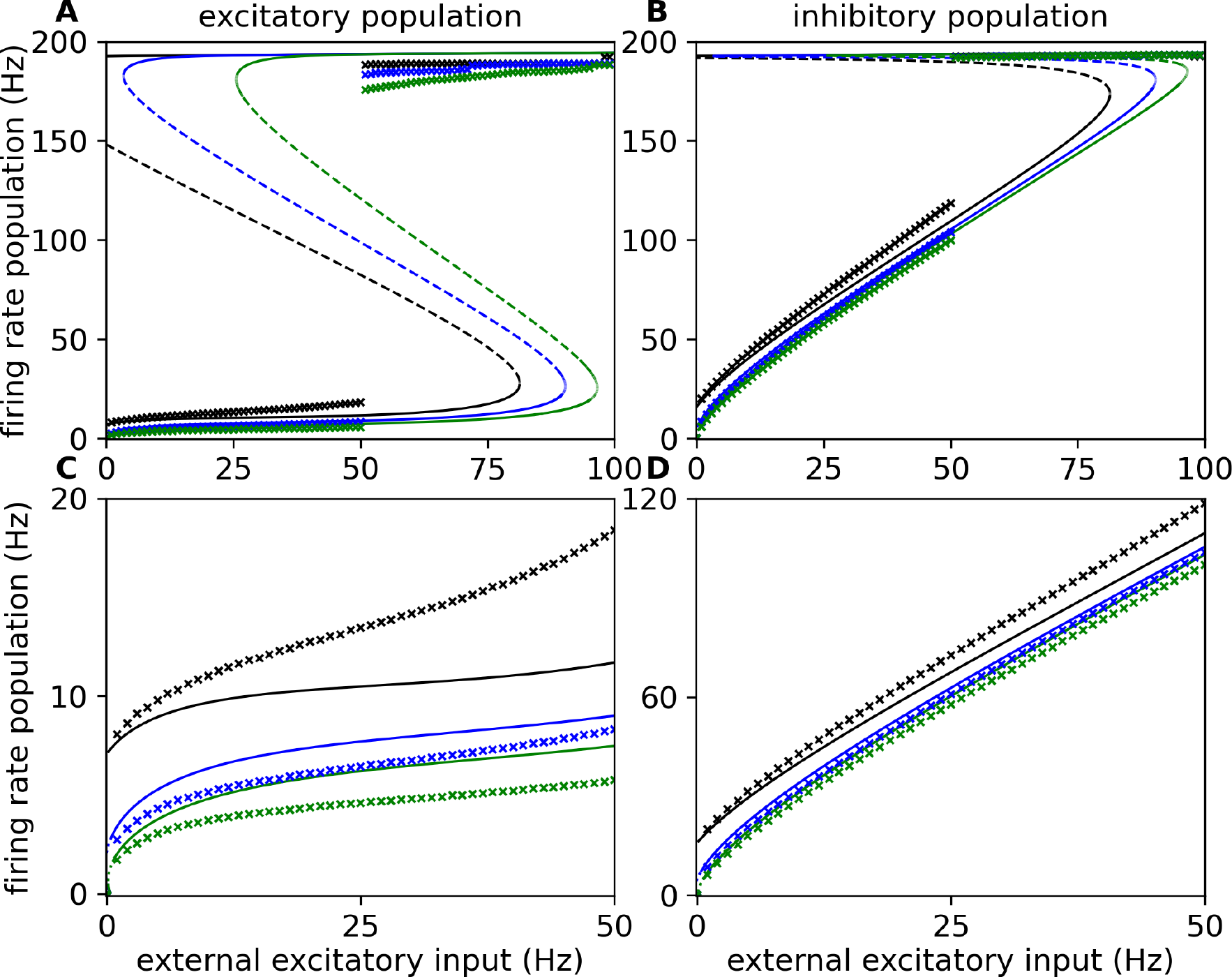
Impact of spike-triggered adaptation on network dynamics. The figure shows the bifurcation diagram, which depends on the external excitatory input for three different values of spike-triggered adaptation (black: 0 pA, blue: 30 pA, green: 60 pA). The three curves represent the stability of the fixed-point (a continuous line indicates a stable fixed-point, and a dashed line represents an unstable fixed-point). The crosses on the diagram are an approximation of the stable fixed-point obtained through simulations. The panels **A** and **B** correspond respectively to the diagrams for the excitatory and inhibitory population. Panels **C** and **D** diagrams are a specific focus on the region of interest for inputs between 0 and 50 Hz

As above, figure 3 confirms that the MF captures the qualitative behavior of the network. However, the MF model overestimates the mean firing rate when it is higher than 150 Hz. For the mean firing rate under 150 Hz, the mean-field underestimates or overestimates depending on the value of the spike-triggered adaptation, *b*. The difference is non-linear with the increase of the external excitatory input. This may be reduced by fitting the population’s transfer function with the network’s steady-state in a specific range of values. However, the structure remains qualitatively the same: a stable fixed-point at the zero input (giving a firing rate = 0) that soon becomes an unstable fixed-point going through a Hopf bifurcation for negative or near-by zero values depending on the level of adaptation *b*. After another Hopf bifurcation, it becomes stable again, entering the main region of interest of the MF where the corresponding spiking network exhibit asynchronous irregular dynamics and where the range of firing rate observed correspond to what can be observed experimentally in the cortex [Ison et al., 2011, Roxin et al., 2011]. It remains stable for a long range of input (≈ 80Hz without adaption and *>* 100Hz with adaptation *b* = 30*pA* or *b* = 60*pA*). A bistability exists, with a higher fixed-point around 200Hz. The supplementary figures 2, 3 and 4 provide the four additional dimensions of the bifurcation diagram. They also demonstrate that the MF does not capture quantitatively the variances and covariance. This may be linked to the size of the windows (see 5 and 6).

Our previous method for estimating the steady-state of the network is limited to one state, and it cannot provide evidence of the existence of bi-stability. By a reduction every 10 seconds of 1Hz of the external input of the Poisson Generator for a network in a steady-state around 200 Hz, we estimate the range of existence of the bi-stability for an external firing rate under 51 Hz. Similarly, by an increase every 10 seconds of 1Hz of the external input of a steady-state network under 150 Hz, we estimate the existence of bi-stability for an external firing rate over 48 Hz. The supplementary figures 7 and 8 show that the MF overestimates the bi-stability’s existence (for details see supplementary note 1). As well as previously, the comparison was also limited to a specific network and one realization of noise. To examine the generalization of the MF, the estimation of the steady-state of the network was done thirty times. The result demonstrates a low variability of the steady-state except when the external input is closed to 50Hz, around the shift of attractor (see supplementary figure 1).

The crucial aspect for the accuracy of the MF AdEx is the fitted transfer function that, in the equations, specifies the dynamics. As detailed in the method section, the final expression is based on the semi-analytical approach, building the link between the single neuron response to inhibitory and excitatory input and the MF. We aim to characterize it in figure 4 provides an analysis of the MF transfer function’s ability to capture the single neuron’s firing rate in response to excitatory input. The top plot (fig. 4A, B) shows the MF transfer function fitted. The middle plot (fig. 4C, D) shows the single neuron firing rate as a function of excitatory input, while the bottom plot (fig. 4E, F) shows the absolute error corresponding to the difference between the MF transfer function and the firing rate of the single neuron. The results show that the transfer function captures the firing rate well, with the maximum error occurring when the inhibitory firing rate is null. For the excitatory neurons transfer function, the maximum error was found to be 40.089 Hz, with an error of 79.769 Hz and a relative error of 0.0056 Hz for the range between 0 Hz and 200 Hz (see supplementary figure 9). It becomes much lower if we only consider the range between 0 Hz and 50 Hz, where the maximum error was found to be 3.407 Hz with an absolute error of 0.1743 Hz and a relative error of 0.0218 Hz. For inhibitory, between 0 Hz and 200 Hz, the maximum error was found to be 34.716, with an absolute error of 57.477 Hz and a relative error of 0.00478 Hz (see supplementary figure 9). If we only consider the range between 0 Hz and 50 Hz, the maximum error was 1.7353 Hz with an absolute error of 0.0688 Hz and a relative error of 0.00687 Hz. In this paper, the fitting of transfer functions includes the effect of the adaptation currents by using data from simulations of neurons with constant negative currents. This was done to reduce the discrepancy between MF and network firing rate when the spike trigger increased, as shown in Di Volo et al. 2019 [di Volo et al., 2019a]. For this set of parameters, the effect of the adaptation does not have a significant impact (under the fitting errors without this additional data), as shown in the supplementary figure 10.

**Figure 4:**
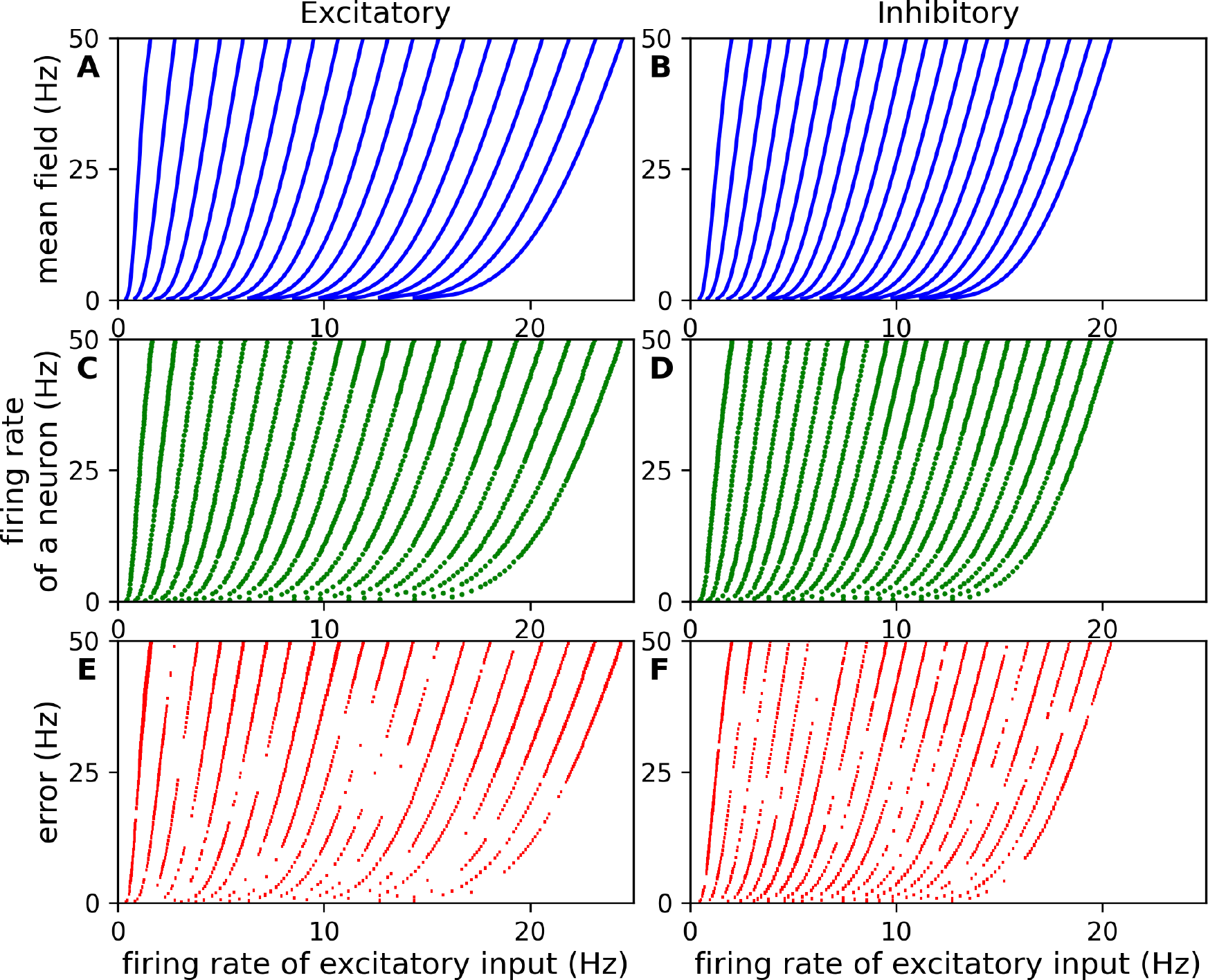
Fitting of Transfer functions. The left part (A, C and E) is dedicated to the excitatory neuron, while the right part (B, D and F) is dedicated to the inhibitory neuron. **A**,**B**: The top graph represents the transfer function of the mean-field fitted to the data. **C, D**: The middle graph shows the data used for the fitting, which is the mean firing rate of 50 independent neurons. **E, F**: The bottom graph displays the error between the transfer function and the data. Each line is associated with a different value of inhibitory firing rate (20 values evenly distributed between 0 Hz and 40 Hz).

Due to the crucial aspect of the transfer function on the MF dynamic, we analyze the sensibility of the precision of the polynomial coefficients of the excitatory transfer function (for details see supplementary note 1). The sensibility analysis is the variation of the 10 Hz fixed-point (far from bifurcation) depending on the number of significant digits of the polynomial coefficients. For an accuracy of 10Hz, it requires at least one significant digit. For an accuracy of 1 Hz, it requires at least three significant digits. And, for an accuracy of 0.1 Hz, it requires at least four significant digits. However, the more the network dynamics is closed to a bifurcation point, the more sensitive the stability of fixed to the precision of the coefficients. Furthermore, for some specific states of the MF, the mean firing rate becomes negative (see supplementary figure 11 and supplementary note 1). Consequently, the MF becomes undefined because the transfer function is not defined for negative firing rates, as expected, negative firing rates are unrealistic cases.

However, another possible source of differences between the network and the MF observed in figure 3 is that the network may not stay in an asynchronous irregular regime while the input increases. While investigating the MF, we did not explore the network behaviors in more detail, which would require an entirely separate study.

In conclusion, the MF captures qualitatively well the different network steady-states. Still, some quantitative differences are observed and may be improved by a better fitting of the transfer function on a specific region of interest for a particular usage of this model.

### 3.2 Dynamical comparison

This part of the analysis is the dynamical response of the MF to non-constant inputs. Indeed, this model is used to constitute a network of MF to model a cortical region [Zerlaut and Destexhe, 2017] or whole brain modeling [Goldman et al., 2020]. In such simulations, it is not anymore the steady-state of the network but the response to varying inputs that is of interest. Thus, the network may not exhibit the asynchronous irregular regime and/or the condition for the Markovianity assumption under non-constant inputs. We tested for different sinusoidal variations of the external Poissonian inputs.

Figure 5 shows the range of validity of the hypothesis of the MF. Indeed, one of the important assumptions to avoid underestimation is the presence of only a single spike of each neuron within the time windows T (see [El Boustani and Destexhe, 2009]). The red line corresponds to the network’s auto-correlation time equals the minimum time between two spikes. The top row of the figure describes the validity range for low amplitude and/or low frequency. It shows that the MF hypothesis is valid for a wide range of low frequency inputs but breaks down for high frequency or/and high amplitude inputs.

**Figure 5:**
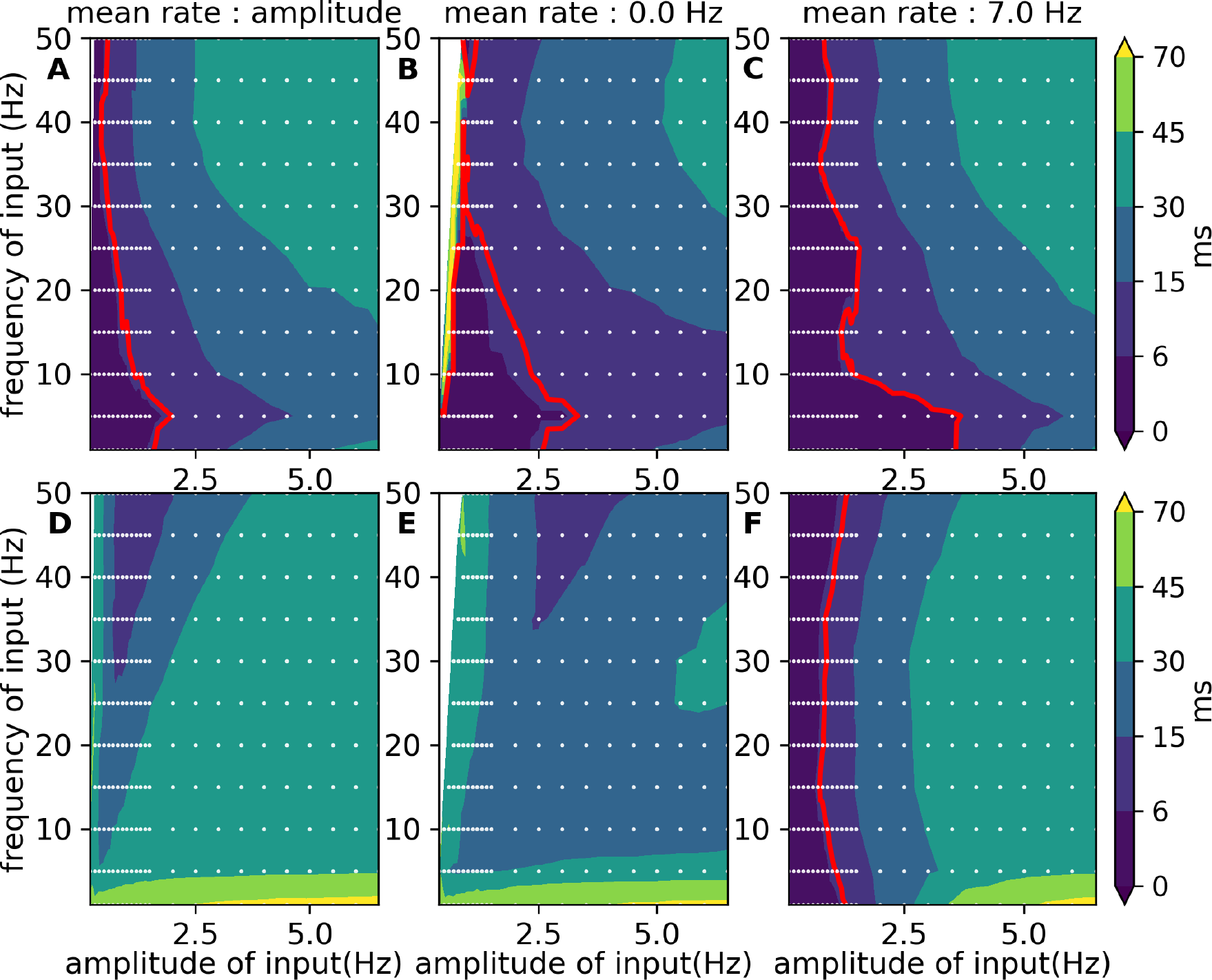
Time scale of the excitatory population in the presence of a sinusoidal Poisson generator. Each graphic represents the timescale of the excitatory population in the presence of stimulation by a sinusoidal Poisson generator mimicking the connection with another population. The red line represents the minimum time between two spikes equals the timescale of the network. The neurons do not have any adaptation for the top graphics (A, B and C), while for the bottom graphics (D, E, F), the value of their spike-triggered adaptation is 60.0 pA. The difference between each column is the mean rate of the sinusoidal Poisson generator. **A, D**: The mean rate equals the amplitude of the oscillation. **B, E**: The mean rate is fixed at 0.0 Hz. **C, F**: The mean rate is fixed at 7.0 Hz, respectively.

On the other hand, the bottom row of the figure shows the effect of adaptation (b=60.0pA) on the range of validity. It shows a validity range only when the average input frequency is 7.0 Hz. In other words, the MF hypothesis, while receiving non-constant input, is valid only for a narrow range of input values when the system has adaptation.

Out of the MF assumptions, we compare the dynamics of the MF and the network in response to a sinusoidal input. Figure 6 shows that the Phase Locking Values between the input signal and the network or MF is close to 1.0, i.e. synchronized. The only exception is when the network receives a signal with low amplitude or low frequency; this can come from the variability of the input of the inhomogeneous Poisson generator. However, the phase shift analysis gives a clear difference between the network and the MF. The MF’s phase shift is more negative than the networks and is in anti-phase with the input for high amplitude and high firing rate (yellow area).

**Figure 6:**
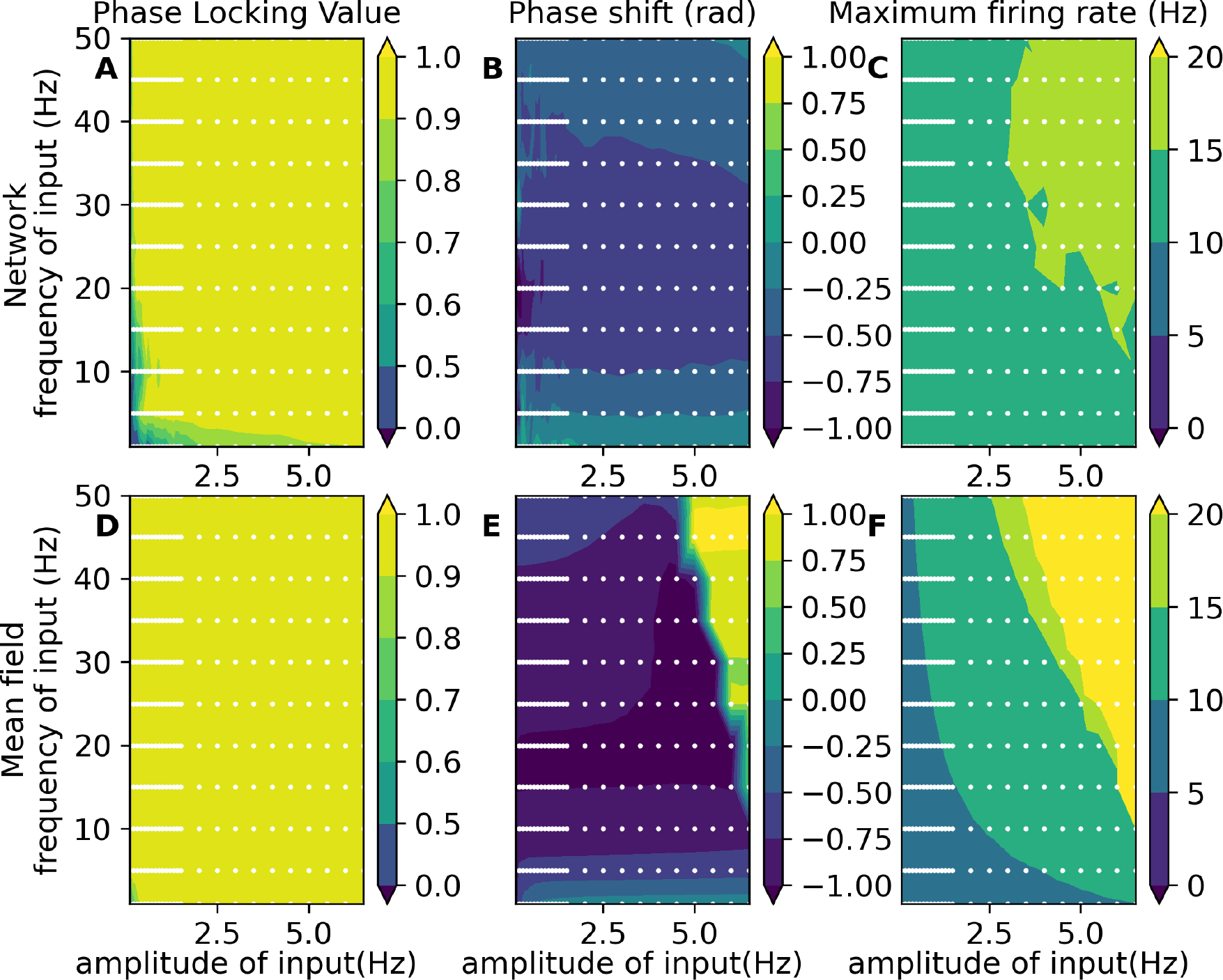
Comparison between the mean-field and the spiking network in the presence of oscillatory input with a mean rate of 7Hz. The graphics **A, B** and **C** represent the measure of the excitatory population in the spiking neural network, and the **D, E** and **F** graphics illustrate the measure of the excitatory mean firing rate of the mean-field. The three measures displayed here are the phase locking values in rad (A and D), the phase shift (B and E) between the oscillatory input and the mean firing rate in rad, and the excitatory population’s maximum firing rate in Hz(C and F).

This area also corresponds to a maximum firing close to 200Hz. On the contrary, the network has a maximum firing rate in the range of 10 Hz to 20 Hz. This means that the network is more attracted around the center fixed-point than the MF, i.e. has a tendency to converge toward this corresponding attractor. Including adaptation does not change these observations but increases the mismatch already noted (see supplementary figure 12). Nevertheless, the mismatches become more significant if the sinusoidal input is centered around 0.0 Hz. As the mean firing rate does not attract to the center fixed-point, it creates a difference in the phase locking values and phase shift (see supplementary figure 13 and 14). The adaptation pushes the network in anti-phase, reducing the discrepancy between the MF and the network for the phase shift. However, the other observation remains the same.

## 4 Discussion

Many different approaches are possible to capture brain dynamics at the mesoscopic scale [Bandyopadhyay et al., 2021, Brunel and Hakim, 2015, Cakan and Obermayer, 2021, El Boustani and Destexhe, 2009, Huang and Lin, 2018, Montbrió et al., 2015, Nykamp and Tranchina, 2000, Schmutz et al., 2020, Schwalger and Chizhov, 2019, Wilson and Cowan, 1972], and each of these approaches is relevant to capture specific features of the dynamics of spiking networks. Some specificities of the mean-field methods studied here are the possibility of considering the sparse connectivity, conductance-based interactions and the finite size of the populations. This method has been applied considering different single neuron models [Alexandersen et al., 2023, Carlu et al., 2020, di Volo et al., 2019b,El Boustani and Destexhe, 2009], but the most used to build large-scale model is based on the AdEx model. Thus, we aim to offer relevant information and characterization approaches about this specific MF model.

The present characterization of the AdEx mean-field (MF) indicates that the MF provides good qualitative insights into the complex behavior of networks of the adaptive exponential integrate and fire neurons with conductance synapses. The approximation of the MF is tightly connected to its transfer function and, in particular, to the precision of the phenomenological threshold. However, this mean-field approximation does not capture the dynamical behaviors of the network for oscillatory inputs.

The bifurcation analysis of the MF reveals the co-existence of multiple fixed-point depending on the external input. In this work, we validated the existence of the co-existence of multi-steady-state in spiking neural networks. Additionally, the MF captures fixed-points around 200 Hz despite the non-validity of the MF’s assumption that the network regimes are asynchronous irregular regimes. However, the quantitative comparison of the fixed-points and the steady-state present a mismatch which is accentuated by the increase of the spike trigger adaptation of the excitatory neurons. This mismatch has multiple causes, such as the size of the windows for the measures, the approximation of the transfer function or/and the degree of network synchronization.

To evaluate the influence of these causes, we quantify the approximation of the transfer function. This quantification informs us about the errors which are not uniform and increase with the output firing rate of the neuron. However, the approximation can be improved if the input and output firing rate range is reduced. Additionally, sensitive analysis of the polynomial coefficients of the phenomenological threshold of the transfer function shows that reducing the imprecision at 0.1 Hz, requires at least 4 significant digits.

In the presence of oscillatory input, we check the co-validity of two hypotheses: the time constant of the MF is high enough for considering that the network is memory-less and low enough for the neurons not to generate more than one spike during this interval. We found that these two hypotheses are valid simultaneously for low amplitudes of the oscillatory input and depend on the average input. Additionally, adding the adaptation current can create cases where these two hypotheses are exclusive because the adaptation current creates a slow fluctuation of mean firing rate. Despite the validity or not of these two hypotheses, the dynamicscomparison between the MF and the network presents a good match for synchronization with the input but a mismatch in mean firing rate, like previously, and a mismatch in the phase shift with the input. These mismatches are variable with the average input and are quite insensitive to the adaptation current. The possible cause of these mismatches is the adiabatic approximation of the MF methods [El Boustani and Destexhe, 2009].

This study is limited to a specific set of parameters and mostly to a specific network connectivity. However, the realization of thirty other networks provides a low variance except around the shift of attractor. Additionally, the qualitative dynamics are similar to that presented in Di Volo et al. 2019 [di Volo et al., 2019a]. The analyses do not provide exhaustive confirmation of all MF’s hypotheses, such as the Gaussian distribution of the voltage membrane, the adiabatic process, and the input spike trains of each neuron are equivalent to a Poisson generator. It explains the partial conclusion of the study but provides a characterization to help the interpretation of MF simulation and clarify the link between the MF and the network. Additionally, we do not provide a study of the different versions of the phenomenological threshold [di Volo et al., 2019a, Zerlaut and Destexhe, 2017, Zerlaut et al., 2018] which can reduce the MF’s approximation. For example, the paper of Zerlaut et al. 2018 [Zerlaut et al., 2018] seems to better approximate the MF because the difference in phase shift is smaller. Still, it was realized with a different transfer function and simulator. Indeed, the implementation of the Poisson generator may differ between simulating environments such as between NEST [Diesmann et al., 2002] and Brian2 [Stimberg et al., 2019], which can impact the network regimes, particularly for high frequency. The good practice will be to perform a characterization of the MF for each specific application. The MF provides a good insight into the network for steady-state. However, the quantitative comparison provides a mismatch between the mean-field and the network, especially with dynamical inputs. Consequently, the usage in the network of mean-field does not ensure its correspondence with the spiking neural network. It requires additional validation for interpreting the MF as a substitution of it.

Such an approach, through a transfer function, can be applied to obtain a mean-field model of very complex neuron models [Alexandersen et al., 2023, Carlu et al., 2020, Zerlaut and Destexhe, 2017]. Future work could be to do a systematic characterization of these other mean-field approximations.

In conclusion, we have characterized here the different dynamical behaviors of the AdEx mean-field model and provided the corresponding bifurcation diagrams. The availability of such diagrams is very useful when searching for specific states of the system, so we anticipate that this information will be highly useful, not only to understand the behavior of these MF models but also to better interpret simulations of large-scale dynamics using networks of synaptically-connected MF models at mesoscale [Zerlaut et al., 2018] or whole-brain scale [Goldman et al., 2020,Goldman et al., 2023].

## Acknowledgments

We thank Mallory Carlu, Matteo di Volo, and Jennifer Goldman for stimulating discussions. This research has received funding from CNRS and the European Union’s Horizon 2020 Framework Programme for Research and Innovation under the Specific Grant Agreement No. 945539 (Human Brain Project SGA3) and Specific Grant Agreement No. 785907 (Human Brain Project SGA2).

## Author contributions

Conceptualization: LK, DD, AD, VJ Formal Analysis: LK Funding Acquisition: VJ, AD Methodology: LK, DD Project Administration: VJ, AD Resources: VJ, AD Software: LK Investigation: LK, DD Visualization: LK, DD Supervision: DD, AD, VJ Writing-original draft: LK, DD Writing-review & editing: LK, DD, AD, VJ

## Data and materials availability

The code for the simulation and analysis of this paper is freely available under v2 Apache license at https://github.com/lionelkusch/compare_zerlaut/tree/paper.

Each exploration’s analysis results are registered in databases available on Zenedo (http://dx.doi.org/10.5281/zenodo.8070791) with the associate code. Supplementary Note 2 describes their contents.

