## Supplementary Material for "Dynamics and bifurcation structure of a mean-field model of adaptive exponential integrate-and-fire networks"

| | |
|---|---|
| Supplementary Note 1 | Additional analysis |
| Supplementary Note 2 | Data available |
| Supplementary Note 3 | supplementary results of spiking neural network |
| Supplementary Note 4 | Supplementary results of mean-field |
| Supplementary Table 1 | Description of the network simulation simulation |
| Supplementary Table 2 | Description of the mean field simulation |
| Supplementary Figure 1 | Variability of fixed-point from network simulation with and without adaptation |
| Supplementary Figure 2 | Bifurcation diagram of external excitatory input for spike-trigger adaptation equals 0.0 pA |
| Supplementary Figure 3 | Bifurcation diagram of external excitatory input for spike-trigger adaptation equals 30.0 pA |
| Supplementary Figure 4 | Bifurcation diagram of external excitatory input for spike-trigger adaptation equals 60.0 pA |
| Supplementary Figure 5 | Variability of the measure of the firing rate |
| Supplementary Figure 6 | Variability of the measure of the firing rate |
| Supplementary Figure 7 | Estimation and comparison of the high fixed-point |
| Supplementary Figure 8 | Estimation and comparison of the low fixed-point |
| Supplementary Figure 9 | Fitting of Transfer functions full |
| Supplementary Figure 10 | Estimation of error of transfer function for neurons in the presence of negative current |
| Supplementary Figure 11 | Negative derivative of the transfer function for zeros firing rate |
| Supplementary Figure 12 | Comparison between the mean-field and the spiking network in the presence of oscillatory input with a mean rate of 7Hz and adaptation (b = 60.0pA) |
| Supplementary Figure 13 | Comparison between the mean-field and the spiking network in the presence of oscillatory input with a mean rate of 0Hz |
| Supplementary Figure 14 | Comparison between the mean-field and the spiking network in the presence of oscillatory input with a mean rate of 0Hz and adaptation (b = 60.0pA) |

### 1 Supplementary Note 1: Additional analysis

Additional analysis was performed to complete the main analysis

#### 1.1 Variability of the measure of the firing rate

The spiking network with a Poisson generator does not have a constant firing rate over time. Measuring the firing rate variability is crucial for comparing with a mean-field because it quantifies the tolerable range of difference with the mean-field. The variability of the mean and standard deviation of the firing rate is estimated with the measure's distribution of 5000 values taken on 30 s of simulation after a transient of 10 s. The only parameter for calculating the firing rate from a spike train using a rectangular window is the window's size. We analyze the influence of the window's size using a log scale between 0.2 ms and 3489.9 ms.

#### 1.2 Estimation of coexistence of fixed-points

A first parameter exploration of the external input gives the values where the network is switching between a fixed-point around 50 Hz to a fixed-point around 200 Hz of excitatory firing rate. From a value of external firing close to this shift, a longer simulation is run where the external input is reduced or increased by 1Hz every 10s. The variation of the mean firing rate estimates the range of coexistence of fixed-points by considering that 10 s is long enough to demonstrate the existence of a steady-state.

#### 1.3 Sensitive analysis of polynomial coefficients of the phenomenal threshold of the transfer function

The sensitivity analysis in this study quantifies the variation of a fixed-point depending on a number of significant digits of the coefficients. We choose to quantify the variability of the fixed-point for $b = 0$ and for an excitatory firing rate equal 10 Hz. A numerical continuation of this fixed-point is performed for each coefficient. From these numerical continuations, we quantify the variation of the polynomial coefficients for the variability of 0.1Hz, 1Hz and 10Hz of the firing rate of the fixed-point. By taking the higher variability of coefficients, we find the minimal number of digits for this three precision of the fixed-point. However, this analysis does not consider the variance due to the variation of the combination of two or more coefficients.

#### 1.4 Existence of negative derivation for null firing rate

The transfer function is not defined for a negative firing rate and the mean-field should not define negative derivation for a null firing rate. However, for specific value of the mean-field is not the case. For identifying these values, we assume that the firing rate of a population is null but also its variance and the covariance. By consequence, the first order equation become:

$$0 < T\,\frac{\partial \nu_e}{\partial t} \tag{1}$$

$$0 < (\mathcal{F}_e - \nu_e) + \frac{1}{2}\,c_{ee}\,\frac{\partial^2 \mathcal{F}_e}{\partial \nu_e \partial \nu_e} + \frac{1}{2}\,c_{ei}\,\frac{\partial^2 \mathcal{F}_e}{\partial \nu_e \partial \nu_i} + \frac{1}{2}\,c_{ie}\,\frac{\partial^2 \mathcal{F}_e}{\partial \nu_i \partial \nu_e} + \frac{1}{2}\,c_{ii}\,\frac{\partial^2 \mathcal{F}_e}{\partial \nu_i \partial \nu_i} \tag{2}$$

$$0 < (\mathcal{F}_e - 0.) + \frac{1}{2}\,0.\,\frac{\partial^2 \mathcal{F}_e}{\partial \nu_e \partial \nu_e} + \frac{1}{2}\,0.\,\frac{\partial^2 \mathcal{F}_e}{\partial \nu_e \partial \nu_i} + \frac{1}{2}\,0.\,\frac{\partial^2 \mathcal{F}_e}{\partial \nu_i \partial \nu_e} + \frac{1}{2}\,c_{ii}\,\frac{\partial^2 \mathcal{F}_e}{\partial \nu_i \partial \nu_i} \tag{3}$$

$$0 < \mathcal{F}_e + \frac{1}{2}\,c_{ii}\,\frac{\partial^2 \mathcal{F}_e}{\partial \nu_i \partial \nu_i} \tag{4}$$

$$c_{ii} < 2\frac{\mathcal{F}_e}{\frac{\partial^2 \mathcal{F}_e}{\partial \nu_i \partial \nu_i}} \tag{5}$$

The supplementary figure 11 displays the minimal positive value of the covariance where this condition is respected.

#### 2 Supplementary Note 2: Data available

In the Zenodo repository, the data from simulations and analyses are available. The available bifurcation analysis results from the numerical continuation of the mean-field for all different values of adaptation spike trigger $b$ (0., 30., 60.). However, the bifurcations for the sensitivity analysis are not available. The repository contains the result of the negative derivative for a null firing rate (see supplementary figure 11) for excitatory and inhibitory transfer functions. It also contains the data for fitting the transfer function and the result of the fit. The only available spike trains are the output of the short simulation with an external input rate of 10, 60 and 80 Hz used in the figure 1. The only additional data not saved in a database is the firing rate of the steady-state of the analysis of the coexistence of fixed-point for each value of b (see supplementary figure 7 and 8). The Zenodo repository contains multiple databases: one database contains the analysis of the spike trains of the steady-state for different external input firing rates for each seed (see supplementary figure 1), one database for each long simulation used in the analysis of the variability of firing rate (see supplementary figure 5 and 6), one database for each parameter exploration of the oscillatory input parameters of neural network (see figure 5) and a unique database for the parameters exploration of oscillatory input parameters for the mean-field (see figure 6).

### 3 Supplementary Note 3: Supplementary results of spiking neural network

#### 3.1 Firing rate and associated measures

The firing rate is the number of generated spikes by a neuron during 1 s. One measure of population synchronization quantifies the fluctuation of the population mean firing rate (for windows of 0.1 and 1 ms) with the coefficient of variation [4]. Additionally, one estimation of neural population frequency uses the dominant frequency and its associated power of the population mean firing rate (for a window of 1 ms) using Welch's method [16] from the elephant package [5]. (Databases record average ('rates_average'), standard deviation ('rates_std'), minimum ('rate_min') and maximum ('rate_max') mean firing rates; for each (0.1 ms and 1 ms) the coefficient of variation ('cvs_IFR_0_1ms' and 'cvs_IFR_1ms'); and for a window of 1 ms, the dominant frequency and its power('frequency_hist_1_freq' and 'frequency_hist_1_val').

#### 3.2 Inter-spiking interval and associated measures

The inter-spiking interval is the time between 2 spikes. The distribution of this interval characterizes the regularity or irregularity of neuron spike generation [4], which is quantified by two measures: coefficient of variation of firing rate [4] and local variation [15][14]. These measures are provided by Elephant python package [5]. These measures are determined for viable estimation by considering only neurons with more than 20 spikes in 10 s for analysis. (Databases record average ('ISI_average'), standard deviation('ISI_std'), minimum ('ISI_min'), and maximum ('ISI_max') inter-spiking intervals; the average and standard deviation over the population of the regularity measures ('cvs_ISI_average', 'cvs_ISI_std', 'lvs_ISI_average' and 'lvs_ISI_std'); and the percentage of neurons with more than 20 spikes ('percentage').

#### 3.3 Phase and associated measures

Kuramoto proposed a model for describing and analyzing synchronization of coupled intrinsic oscillators [11], based on the following equation:

$$\frac{\partial \theta_i}{\partial t} = \omega_i + \frac{K}{N}\sum_{j=1}^{N} \sin(\theta_j - \theta_i),\ i = 1...N, \tag{6}$$

where the system is composed of N limit-cycle oscillators, with phases $\theta_i$ and coupling constant K.

The phase of a neuron can be defined by approximating the voltage membrane as an intrinsic oscillator or by approximating the phase using a constant velocity between 2 spikes, similar to Borges et al. 2017 [2]. This study used the second approximation described in the following equation:

$$\psi(t) = 2\pi m + 2\pi \frac{t - t_m}{t_{m+1} - t_m}, \forall t \in [t_m, t_{m+1}] \tag{7}$$

where $t_m$ is the m-th spike of the neuron.
One measure of synchronization is the estimation of Kuramoto order-parameter of a population, i.e. the absolute values of the mean phase. Additionally, one estimation of neural population frequency uses the dominant frequency and its associated power of the mean phase (for a window of 0.1 ms) using Welch's method [16] from the elephant package [5]. (Databases record average ('synch_Rs_average'), standard deviation ('synch_Rs_std'), minimum ('synch_Rs_min') and maximum ('synch_Rs_max') of the Kuramoto order-parameter; the start time and end time of the mean phase estimation ('synch_Rs_times_init' and 'synch_Rs_times_end'); and for a window of 0.1 ms, the dominant frequency and its power ('frequency_phase_freq' and 'frequency_phase_val').)

#### 3.4 Bursts and associated measures

Bursts are consecutive spikes with inter-spike intervals of less than 10 ms. Additionally, the regularity of bursts is analyzed similarly to inter-spike intervals. For neurons with more than 20 bursts, the regularity of the interval of the beginning time of bursts and also their ending time is quantified with the coefficient of variation and the local variation. (Databases record average ('burst_count_average', 'burst_nb_average' and 'burst_rate_average'), standard deviation ('burst_count_std', 'burst_nb_std' and 'burst_rate_std'), maximum ('burst_count_max', 'burst_nb_max' and 'burst_rate_max') and minimum ('burst_count_min', 'burst_nb_min' and 'burst_rate_min') of a number of spikes in the sequence of burst, number of burst by neurons during the analysis and mean bursting rates; the percentage of neurons which have one burst ('percentage_burst'); average ("burst_interval_average'), standard deviation ('burst_interval_std'), minimum ('burst_interval_min'), maximum ('burst_interval_max'); and for neurons with more than 20 bursts, the average and the standard deviation of the coefficient of variation ('burst_cv_begin_average', 'burst_cv_end_average', 'burst_cv_begin_std', 'burst_cv_end_std') and local variation ('burst_lv_begin_std', 'burst_lv_end_std', 'burst_lv_begin_average', 'burst_lv_end_average') of interval of the beginning time of bursts and also their ending time.

#### 3.5 Relation with input firing rate

The mean phase difference between the oscillatory input and the oscillatory histogram of the spiking neural network:

$$meanphase = \frac{1}{N} \sum_{n=1}^{N} \exp\left(j\left(\theta_{input}(t,n) - \theta_{network}(t,n)\right)\right) \tag{8}$$

where $\theta_{input}$ is the analytic signal given by the Hilbert transformation of the sinusoidal input, and $\theta_{network}$ is the analytic signal provided by the Hilbert transformation of the network oscillation signal. The absolute value of the *meanphase* corresponds to the phase locking value as defined by Lachaux et al. 1999 [12]. The angle of the *meanphase* corresponds to the phase shift[9]. This measure is calculated for a histogram with a bin of 0.1 ms, 1.0 ms and 5 ms and a smoothed histogram with an average sliding window of 5 ms. Additionally, the time scale of the network is only analyzed for networks with oscillatory input. The time scale is quantified using the auto-correlation function of elephant package [5]. The function is the implementation of the method proposed by Wieland et al. 2015 [17]:

$$\tau_{\text{corr}} = \int_{-\tau_{\max}}^{\tau_{\max}} \left[\frac{\hat{C}(\tau)}{\hat{C}(0)}\right]^2 d\tau \tag{9}$$

where $\hat{C}(\tau) = \langle x(t)x(t-\tau)\rangle \langle x\rangle^2 - \langle x\rangle \delta(\tau)$ with $x(t) = \sum_{n=1}^{N} \delta(t - t_n)$ for $t_n$ is the n-th spike time of the spike trains.
(Databases with oscillatory input record phase locking value ('PLV_0_1ms', 'PLV_1ms', 'PLV_5ms', 'PLV_w5ms'), phase shift ('PLV_angle_0_1ms', 'PLV_angle_1ms', 'PLV_angle_5ms', 'PLV_angle_w5ms' and 'MeanPhaseShift_0_1ms', 'MeanPhaseShift_1ms', 'MeanPhaseShift_5ms', 'MeanPhaseShift_w5ms'), the coefficient of variation of the histogram ('cvs_IFR_0_1ms', 'cvs_IFR_1ms', 'cvs_IFR_5ms', 'cvs_IFR_w5ms'), maximum of firing rate (max_IFR_0_1ms, max_IFR_1ms, max_IFR_5ms, max_IFR_w5ms) for a bin of 0.1 ms, 1 ms, 5 ms and smoothing with windows of 5 ms.)

### 4 Supplementary Note 4: Supplementary results of mean-field

#### 4.1 Firing rate

The firing rate correspond to the variable of the mean-field. This measure variable is quantified, using similar methods that for the histogram of spiking neural network, phase locking value, phase shift and dominant frequency. (Databases with oscillatory input record average ('mean_rates'), standard deviation ('std_rates'), minimum ('min_rates'), maximum ('max_rate') and dominant frequency ('frequency_dom') of firing rates and phase locking value ('PLV_value') and phase shift ('PLV_angle') between the oscillatory input and the firing rate)

### 5 Supplementary Tabular 1: Description of the network simulation simulation

| A1 | NEST : model summary | |
|---|---|---|
| | software | NEST |
| | topology | Erdős - Rényi [8] |
| | population | excitatory and inhibitory |
| | connectivity | random connectivity without self-recurrent connections |
| | neuron model | adaptive exponential leaky integrate and fire neurons, fixed threshold and fixed absolute refractory time |
| | synapse model | conductance-based exponential shape |
| | plasticity | none |
| | input | independent fixed rate Poisson generator spike trains to all neurons |
| | measurement | spike activity |

| A2 | NEST : software | |
|---|---|---|
| | version | alpha 3.0.0 (8f5a5f)[10] |
| | integrator method | 4th order Runge-Kutta-Fehlberg method |
| | integration step | 0.1 ms |
| | number of seeds | 8 (master seed: 0-30 or 46) |
| | simulation time | 2s (example), 5s(normal, transient:1s), 40s(long, transient:10s) or variable(test stability, transient:10s) |

| A3 | NEST : populations | | |
|---|---|---|---|
| | **name** | **elements** | **size** |
| | E | aeif_cond_exp | N_e = (1-$g_{inh}$)N = 8000 |
| | I | aeif_cond_exp | N_i = $g_{inh}$N = 2000 |
| | $P_{ext}$ | Poisson generator | 8000 |
| | $P_{sin\ ext}$ | Sinusoidal Poisson generator | 1 |

| A4 | NEST : neuron model |
|---|---|
| name | aeif |
| type | adaptive exponential leaky integrator and fire [3] with conductance synapse |
| subtreshold dynamics | $$C_m \frac{dV_m}{dt} = - g_L(V_m - E_L) + g_L \Delta_T e^{\frac{V_m - V_{th}}{\Delta_T}} - g_e(t)(V_m - E_{ex}) - g_i(t)(V_m - E_{in}) + I_e - w + I_e$$ $$\tau_w \frac{dw}{dt} = a(V_m - E_L) - w$$ |
| reset condition | for $t^{(f)} = \{t \vert V_m(t) >= V_{peak}\}$ <br>• $V_m([t^{(f)}; t^{(f)} + t_{ref}]) = V_{reset}$ <br>• $w([t^{(f)}]) = w([t^{(f)}]) + b$ |
| | NEST : synapse model |
| name | cond_exp |
| type | post-synaptic conductance in the form of truncated exponentials |
| coupling equation | $$g_e(t) = \sum_{t_j^{(f)}} w_j exp\_trunc(t - t_j, \tau_{syn}) \text{ with } w_j > 0.0$$ $$g_i(t) = \sum_{t_j^{(f)}} w_j exp\_trunc(t - t_j, \tau_{syn}) \text{ with } w_j < 0.0$$ $$exp\_trunc(t, \tau) = e^{1-\frac{t}{\tau}} Heaviside(t)$$ |

| A5 | NEST : neuron model parameters | |
|---|---|---|
| | excitatory neurons | |
| $C_m$ | capacity of the membrane | 200.0 pF |
| $t_{ref}$ | duration of refractory period | 5.0 ms |
| $V_{reset}$ | reset value for $V_m$ after a spike | -55.0 mV |
| $E_L$ | leak reversal potential | -63.0 mV |
| $g_L$ | leak conductance | 10.0 nS |
| $\Delta_T$ | slope factor | 2.0 mV |
| $V_{peak}$ | spike detection threshold | 0.0 mV |
| $a$ | subthreshold adaptation | 0.0 nS |
| $b$ | spike-triggered adaptation | 0.0/30.0/60.0pA |
| $\tau_w$ | adaptation time constant | 500.0 ms |
| $V_{th}$ | spike initiation threshold | -50.0 mV |
| $I_e$ | constant external input current | 0.0 pA |
| $E_{ex}$ | excitatory reversal potential | 0.0 mV |
| $E_{in}$ | inhibitory reversal potential | -80.0 mV |
| $V_m$ | initialization of the voltage membrane | $-65.0 \pm 100.0$ mV |
| $w$ | initialization of adaptation current | $200.0 \pm 200$ pA |
| $g_e$ | initialization of excitatory synapse input | 0.0 pA |
| $g_i$ | initialization of inhibitory synapse input | 0.0 pA |
| | inhibitory neurons | |
| $C_m$ | capacity of the membrane | 200.0 pF |
| $t_{ref}$ | duration of refractory period | 5.0 ms |
| $V_{reset}$ | reset value for $V_m$ after a spike | -65.0 mV |
| $E_L$ | leak reversal potential | -65.0 mV |
| $g_L$ | leak conductance | 10.0 nS |
| $\Delta_T$ | slope factor | 0.5 ms |
| $V_{peak}$ | spike detection threshold | 0.0 mV |
| $a$ | subthreshold adaptation | 0.0 nS |
| $b$ | spike-triggered adaptation | 0.0 pA |
| $\tau_w$ | Adaptation time constant | 1.0 ms |
| $V_{th}$ | spike initiation threshold | -50.0 mV |
| $I_e$ | constant external input current | 0.0 pA |
| $E_{ex}$ | excitatory reversal potential | 0.0 mV |
| $E_{in}$ | inhibitory reversal potential | -80.0 mV |
| $V_m$ | initialization of the voltage membrane | $-65.0 \pm 100.0$ mV |
| $w$ | initialization of adaptation current | $200.0 \pm 200$ pA |
| $g_e$ | initialization of excitatory synapse input | 0.0 pA |
| $g_i$ | initialization of inhibitory synapse input | 0.0 pA |

| A6 | NEST : input |
|---|---|
| Poisson generator | |
| equation | $p(n) = \frac{\lambda^n}{n!} \exp(-\lambda)$ |
| implementation algorithm | Ahrens and Dieter 1982[1] |
| excitatory firing rate $\lambda_{ex}$ | 0-100 Hz |
| Sinusoidal Poisson generator | |
| equation | $p(n) = \frac{(rate + a \sin(f * n))^n}{n!} \exp(-rate + a \sin(f * n))$ |
| amplitude $a$ | 0-50 Hz |
| frequency $f$ | 0-50 Hz |
| average $rate$ | 0-50 Hz |

| A7 | NEST : connectivity | | | |
|---|---|---|---|---|
| parameter synapses | | | | |
| $\tau_{ex}$ | rise time of excitatory synaptic conductance | | | 5.0ms |
| $\tau_{in}$ | rise time of inhibitory synaptic conductance | | | 5.0ms |
| **name** | **source** | **target** | **weights** | **pattern** |
| EE | E | E | $Q_e$=1.5 nS | probability of connections is 0.05 between pairs of neurons. Neurons cannot be directly connected to themselves nor have multiple connections with the same postsynaptic neuron. |
| EI | E | I | $Q_e$=1.5 nS | probability of connections is 0.05 between pairs of neurons. Neurons cannot be directly connected to themselves nor have multiple connections with the same postsynaptic neuron. |
| IE | I | E | $Q_i$=5.0 nS | probability of connections is 0.05 between pairs of neurons. Neurons cannot be directly connected to themselves nor have multiple connections with the same postsynaptic neuron. |
| II | I | I | $Q_i$=5.0 nS | probability of connections is 0.05 between pairs of neurons. Neurons cannot be directly connected to themselves nor have multiple connections with the same postsynaptic neuron. |
| $P_{ext}$E | $P_{ext}$ | E | $Q_e$=1.5 nS | one generator by neuron. |
| $P_{sin\ ext}$E | $P_{sin\ ext}$ | E | $Q_e$=1.5 nS | the generator is connected to a neurons with a probability of 0.05. |

| A8 | NEST : measurement | | |
|---|---|---|---|
| state variable | spike time | precision | 0.1 ms |
| spike activities | raster plot | precision | 0.1 ms |
| | simple moving average | windows size | T (5ms) |

#### 6 Supplementary Tabular 2: Description of the mean field simulation

| B1 | TVB : model summary |
|---|---|
| software | The Virtual Brain [13] |
| neural mass model | mean-field of Ad Ex[7, 6] |
| connectivity | None |
| input | sinusoidal input |
| monitor | raw values |

| B2 | TVB : software |
|---|---|
| version | 2.6 |
| integrator method | Heun integrator |
| integration step | 0.1 ms |
| simulation time | 20.001 s (including a transient of 2.5 s) |

| B3 | TVB : input and monitor | | |
|---|---|---|---|
| input | sinusoidal input | equation | $rate + a\sin(f * t)$ |
| | | amplitude $a$ | 0-50 Hz |
| | | frequency $f$ | 0-50 Hz |
| | | rate $rate$ | 0 or 7 Hz |
| monitor | Raw monitor | state variable | 0.1 ms |

| B4 | TVB : neural mass model |
|---|---|
| name | Mean-Field AdEx[7] |
| type | neural mass model network of adaptive exponential integrate and fire excitatory and inhibitory neurons of second statistical order |
| equation | $T\,\frac{\partial \nu_e}{\partial t} =(\mathcal{F}_e-\nu_e)+\frac{1}{2}\,c_{ee}\,\frac{\partial^2 \mathcal{F}_e}{\partial \nu_e \partial \nu_e}+\frac{1}{2}\,c_{ei}\,\frac{\partial^2 \mathcal{F}_e}{\partial \nu_e \partial \nu_i}+\frac{1}{2}\,c_{ie}\,\frac{\partial^2 \mathcal{F}_e}{\partial \nu_i \partial \nu_e}+\frac{1}{2}\,c_{ii}\,\frac{\partial^2 \mathcal{F}_e}{\partial \nu_i \partial \nu_i}$<br>$T\,\frac{\partial \nu_i}{\partial t} =(\mathcal{F}_i-\nu_i)+\frac{1}{2}\,c_{ee}\,\frac{\partial^2 \mathcal{F}_i}{\partial \nu_e \partial \nu_e}+\frac{1}{2}\,c_{ei}\,\frac{\partial^2 \mathcal{F}_i}{\partial \nu_e \partial \nu_i}+\frac{1}{2}\,c_{ie}\,\frac{\partial^2 \mathcal{F}_i}{\partial \nu_i \partial \nu_e}+\frac{1}{2}\,c_{ii}\,\frac{\partial^2 \mathcal{F}_i}{\partial \nu_i \partial \nu_i}$<br>$T\,\frac{\partial c_{ee}}{\partial t} =(\mathcal{F}_e-\nu_e)\,(\mathcal{F}_e-\nu_e)+c_{ee}\frac{\partial \mathcal{F}_e}{\partial \nu_e}+c_{ee}\frac{\partial \mathcal{F}_e}{\partial \nu_e}+c_{ei}\frac{\partial \mathcal{F}_e}{\partial \nu_i}+c_{ie}\frac{\partial \mathcal{F}_e}{\partial \nu_i}-2c_{ee}$<br>$+\frac{\mathcal{F}_e\,(1/T-\mathcal{F}_e)}{N_e}$<br>$T\,\frac{\partial c_{ei}}{\partial t} =(\mathcal{F}_e-\nu_e)\,(\mathcal{F}_i-\nu_i)+c_{ee}\frac{\partial \mathcal{F}_i}{\partial \nu_e}+c_{ie}\frac{\partial \mathcal{F}_i}{\partial \nu_i}+c_{ei}\frac{\partial \mathcal{F}_e}{\partial \nu_e}+c_{ii}\frac{\partial \mathcal{F}_e}{\partial \nu_i}-2c_{ei}$<br>$T\,\frac{\partial c_{ie}}{\partial t} =(\mathcal{F}_i-\nu_i)\,(\mathcal{F}_e-\nu_e)+c_{ee}\frac{\partial \mathcal{F}_i}{\partial \nu_e}+c_{ie}\frac{\partial \mathcal{F}_i}{\partial \nu_i}+c_{ei}\frac{\partial \mathcal{F}_e}{\partial \nu_e}+c_{ii}\frac{\partial \mathcal{F}_e}{\partial \nu_i}-2c_{ie}$<br>$T\,\frac{\partial c_{ii}}{\partial t} =(\mathcal{F}_i-\nu_i)\,(\mathcal{F}_i-\nu_i)+c_{ie}\frac{\partial \mathcal{F}_i}{\partial \nu_e}+c_{ei}\frac{\partial \mathcal{F}_i}{\partial \nu_e}+c_{ii}\frac{\partial \mathcal{F}_i}{\partial \nu_i}+c_{ii}\frac{\partial \mathcal{F}_i}{\partial \nu_i}-2c_{ii}$<br>$+\frac{\mathcal{F}_i\,(1/T-\mathcal{F}_i)}{N_i}$<br>$\tau_{W_e}\frac{\partial W_e}{\partial t} =-W_e+b_e\nu_e+a_e(\mu_V(\nu_e,\nu_i,W_e)-EL_e)$<br>$\tau_{W_i}\frac{\partial W_i}{\partial t} =-W_i+b_i\nu_i+a_i(\mu_V(\nu_e,\nu_i,W_i)-EL_i)$ |
| transfer function | $\mathcal{F}_e =\mathcal{F}((\nu_e+1e-6),\nu_{ext},\nu_i,W_e)$<br>$\mathcal{F}_i =\mathcal{F}((\nu_e+1e-6),\nu_{ext},\nu_i,W_i)$<br>$\mathcal{F} =\frac{1}{2\,\tau_V}\cdot Erfc(\frac{V^{eff}_{thre}-\mu_V}{\sqrt{2}\,\sigma_V})$<br>$V^{eff}_{thre}(\mu_V,\sigma_V,\tau_V^N =\tau_V\frac{g_L}{Cm})=P'_0+\sum_{x\in\{\mu_V,\sigma_V,\tau_V^N\}}P_x\cdot\left(\frac{x-x^0}{\delta x^0}\right)$<br>$+\sum_{x,y\in\{\mu_V,\sigma_V,\tau_V^N\}^2}P_{xy}\cdot\left(\frac{x-x^0}{\delta x^0}\right)\left(\frac{y-y^0}{\delta y^0}\right)$<br>$\mu_G(\nu_e,\nu_{ext},\nu_i)=((\nu_eK_e+\nu_{ext}K_{ext})\tau_eQ_e)+(\nu_iK_i\tau_iQ_i)+g_L$<br>$\mu_{V_s}(\nu_e,\nu_{ext},\nu_i,W,\mu_G)=\frac{((\nu_eK_e+\nu_{ext}K_{ext})\tau_eQ_e)E_e+(\nu_iK_i\tau_iQ_i)E_i+g_LEL_s-W}{\mu_G}$<br>$\sigma_V(\mu_V,\mu_G)=\sqrt{\sum_{s\in\{e,i\}}K_s\nu_s\frac{\left(\frac{Q_s}{\mu_G}(E_s-\mu_V)\tau_s\right)^2}{2\frac{C_m}{\mu_G}+\tau_s}}$<br>$\tau_V(\mu_V,\mu_G)=\frac{\sum_{s\in\{e,i\}}K_s\nu_s\left(\frac{Q_s}{\mu_G}(E_s-\mu_V)\tau_s\right)^2}{\sum_{s\in\{ex,in\}}K_s\nu_s\frac{\left(\frac{Q_s}{\mu_G}(E_s-\mu_V)\tau_s\right)^2}{2\frac{C_m}{\mu_G}+\tau_s}}$ |

| B5 | TVB : neural mass model parameters | |
|---|---|---|
| $T$ | time resolution of the mean-field | 5.0 ms |
| $C_m$ | capacity of the membrane | 200.0 pF |
| $EL_e$ | leak reversal potential excitatory($E_L$) | -63.0 mV |
| $EL_i$ | leak reversal potential inhibitory($E_L$) | -65.0 mV |
| $g_L$ | leak conductance | 10.0 nS |
| $a_e$ | subthreshold adaptation of excitatory neurons($a$) | 0.0 nS |
| $b_e$ | spike-triggered adaptation of excitatory neurons($b$) | 0.0/30.0/60.0 pA |
| $\tau_{W_e}$ | adaptation time constant of excitatory neurons($\tau_w$) | 500.0 ms |
| $a_i$ | subthreshold adaptation of inhibitory neurons($a$) | 0.0 nS |
| $b_i$ | spike-triggered adaptation inhibitory neurons($b$) | 0.0 pA |
| $\tau_{W_i}$ | adaptation time constant of inhibitory neurons($\tau_w$) | 1.0 ms |
| $E_e$ | excitatory reversal potential($E_{ex}$) | 0.0 mV |
| $\tau_e$ | rise time of excitatory synaptic conductance($\tau_{ex}$) | 5.0 ms |
| $Q_e$ | excitatory quantal conductance | 1.5 nS |
| $E_i$ | inhibitory reversal potential($E_{in}$) | -80.0 mV |
| $\tau_i$ | rise time of inhibitory synaptic conductance($\tau_{in}$) | 5.0 ms |
| $Q_i$ | inhibitory quantal conductance | 5.0 nS |
| $p_{connect}$ | probability of connection | 0.05 |
| $N_{tot}$ | number of total neurons | 10000 |
| $p_i$ | percentage of inhibitory neurons | 0.2 |
| $N_e$ | number of excitatory neurons | $N_{tot}(1-p_i)$ =8000 |
| $N_i$ | number of inhibitory neurons | $N_{tot}p_i$ =2000 |
| $K_e$ | mean number of input excitatory synapses : $N_e p_{connect}$ | 400 |
| $K_i$ | mean number of input inhibitory synapses : $N_i p_{connect}$ | 100 |
| $\nu_{ext}$ | external input | see external input section |
| | initial condition | $\mu_E : 0.Hz; \mu_i : 0.Hz; c_{ee} : 0.; c_{ei} : 0.; c_{ii} : 0.; W_e : 1000.pA; W_i : 0.pA$ |

| | | | | | |
|---|---|---|---|---|---|
| $P_e$ | second order polynomial of the phenomenological threshold for inhibitory neuron in mV | $P_0$ | $P_{\mu_V}$ | $P_{\sigma_V}$ | $P_{\tau_V^N}$ |
| | | -4.923163e-02 | 1.762790e-03 | -7.677835e-04 | -3.824880e-03 |
| | | $P_{\mu_V^2}$ | $P_{\sigma_V^2}$ | $P_{(\tau_V^N)^2}$ | |
| | | 2.356120e-04 | 4.0210098e-03 | 1.812297e-03 | |
| | | $P_{\mu_V \sigma_V}$ | $P_{\mu_V \tau_V^N}$ | $P_{\sigma_V \tau_V^N}$ | |
| | | -3.723180e-05 | 1.929229e-04 | 3.974934e-03 | |
| $P_i$ | second order polynomial of the phenomenological threshold for inhibitory neuron in mV | $P_0$ | $P_{\mu_V}$ | $P_{\sigma_V}$ | $P_{\tau_V^N}$ |
| | | -5.079953e-2 | 2.139835e-3 | -4.646189e-3 | 3.727148e-4 |
| | | $P_{\mu_V^2}$ | $P_{\sigma_V^2}$ | $P_{(\tau_V^N)^2}$ | |
| | | 5.053228e-4 | 1.304294e-3 | -1.073580e-2 | |
| | | $P_{\mu_V \sigma_V}$ | $P_{\mu_V \tau_V^N}$ | $P_{\sigma_V \tau_V^N}$ | |
| | | 1.995937e-3 | 1.932031e-3 | -1.015957e-2 | |

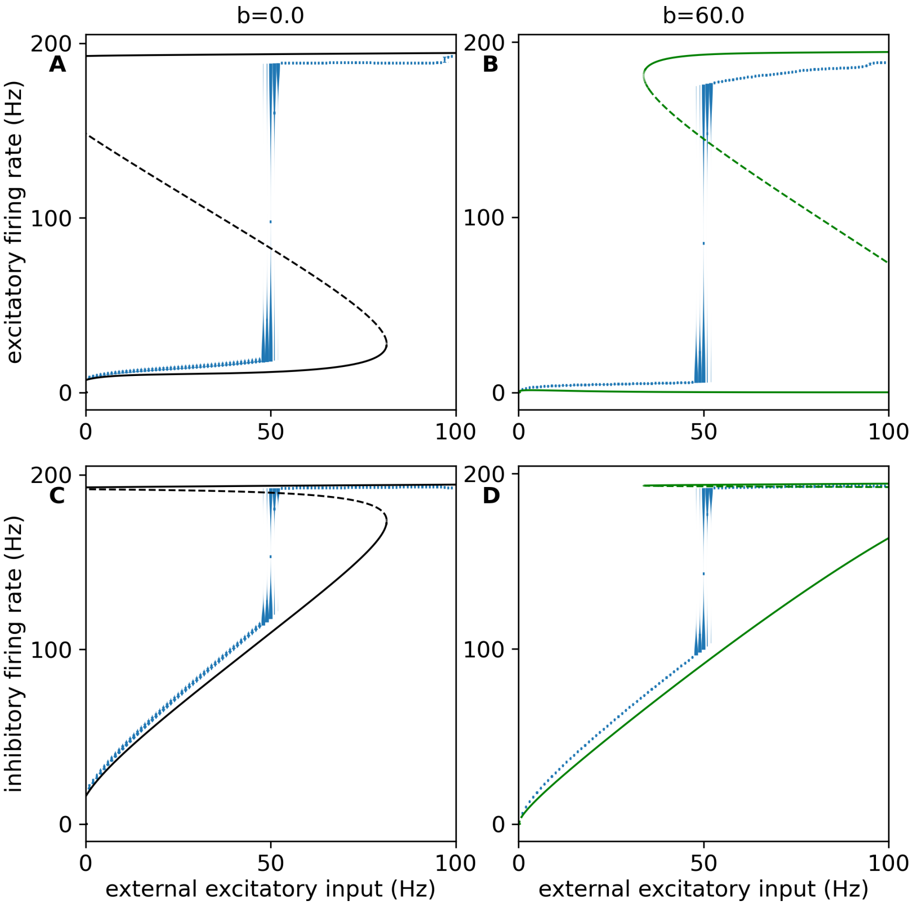


Figure 1: **Variability of fixed-point from network simulation with and without adaptation** Each violin represents the distribution of the mean firing rate of an excitatory or inhibitory population of 4 seconds after 1 second of transient for 30 networks. **A**, **B**: These violins in the top graphics represent the distribution of the estimated fixed-point for the mean firing rate of excitatory dependent on the excitatory external firing rate. **C**, **D**: These violins in the top graphics represent the distribution of the estimated fixed-point for the mean firing rate of excitatory dependent on the excitatory external firing rate. The additional curve represents the bifurcation diagram of the mean-field. The left graphics (A and C) is for cases without adaptation, and the right graphics (B and D) is for cases with spike-trigger adaptation equals 60 pA.

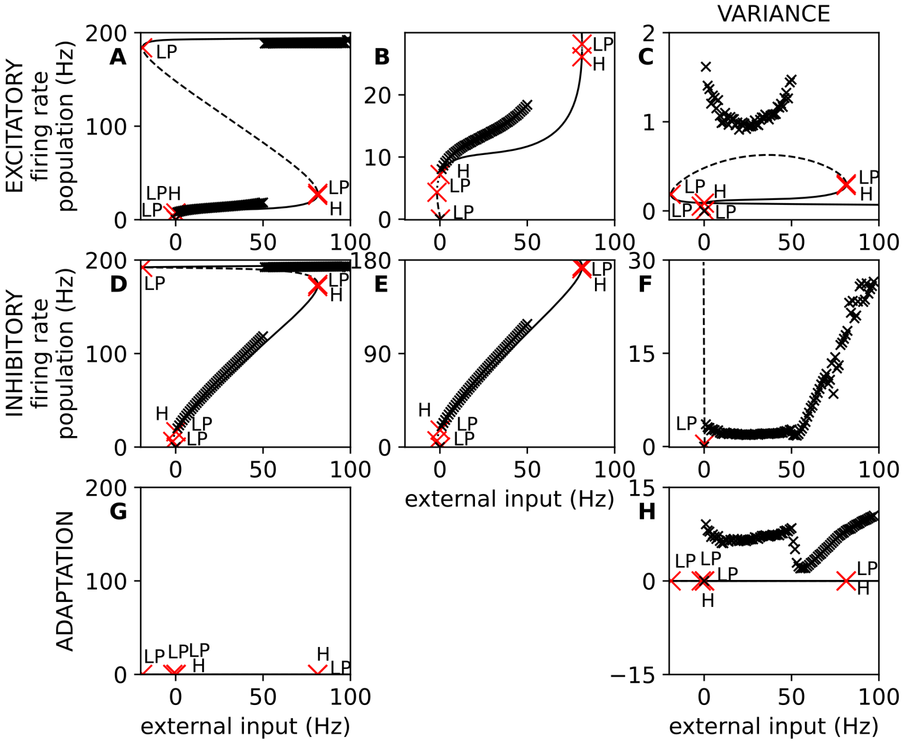


Figure 2: **Bifurcation diagram of external excitatory input for spike-trigger adaptation equals 0.0 pA** Each graphic represents the bifurcation diagram for each variable of the mean-field (**A**: firing rate of excitatory population, **D**: firing rate of inhibitory population, **G**: mean adaptation current, **C**: variance of excitatory firing rate, **F**: variance of inhibitory firing rate, **H**: covariance of the firing rate of the inhibitory and excitatory population). The black curve style represents the stability of the fixed-point (continuous line for stable fix points and dashed lines for unstable fix points). The curve style represents the stability of the fixed-point (continuous line for stable fix points and dashed lines for unstable fix points). The graphics **B** and **E** enlarge the bottom part of their left graphic (A and D). The black crosses are the estimation of the fixed-point using network simulation. The bifurcation points are indicated with a red cross and a letter that means the type of bifurcation (H: Hopf bifurcation and LP: limit point or saddle-node bifurcation).

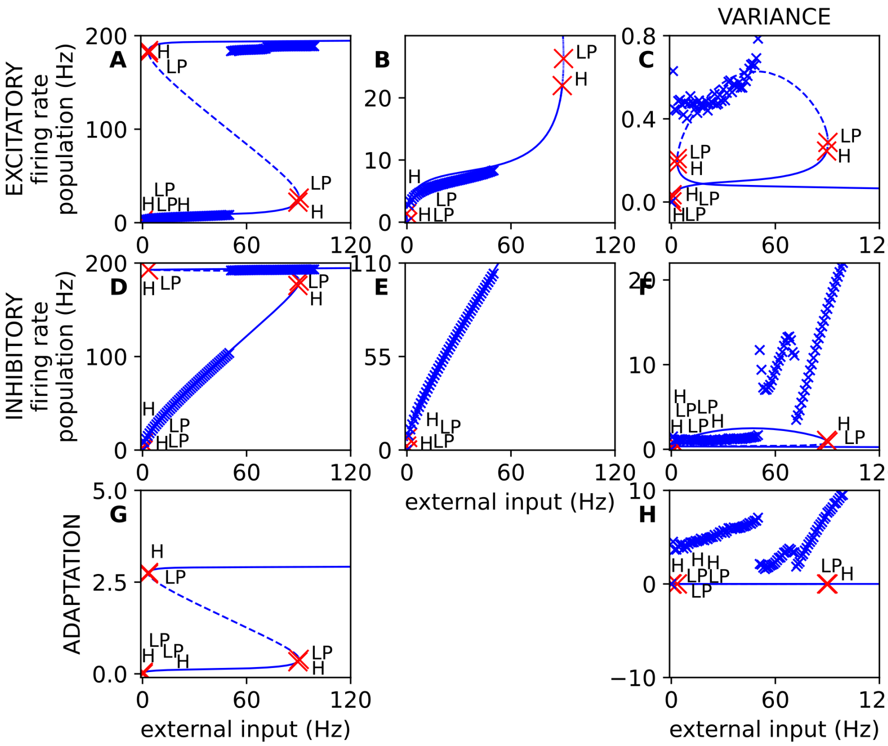


Figure 3: **Bifurcation diagram of external excitatory input for spike-trigger adaptation equals 30.0 pA** Each graphic represents the bifurcation diagram for each variable of the mean-field (**A**: firing rate of excitatory population, **D**: firing rate of inhibitory population, **G**: mean adaptation current, **C**: variance of excitatory firing rate, **F**: variance of inhibitory firing rate, **H**: covariance of the firing rate of the inhibitory and excitatory population). The blue curve style represents the stability of the fixed-point (continuous line for stable fix points and dashed lines for unstable fix points). The curve style represents the stability of the fixed-point (continuous line for stable fix points and dashed lines for unstable fix points). The graphics **B** and **E** enlarge the bottom part of their left graphic (A and D). The blue crosses are the estimation of the fixed-point using network simulation. The bifurcation points are indicated with a red cross and a letter that means the type of bifurcation (H: Hopf bifurcation and LP: limit point or saddle-node bifurcation).

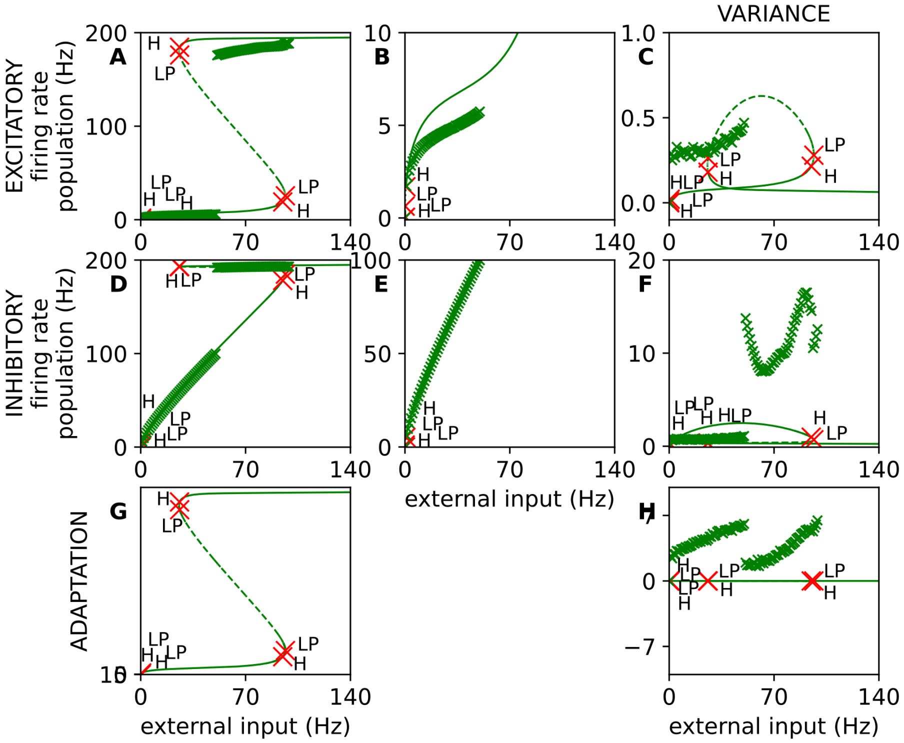


Figure 4: **Bifurcation diagram of external excitatory input for spike-trigger adaptation equals 60.0 pA** Each graphic represents the bifurcation diagram for each variable of the mean-field (**A**: firing rate of excitatory population, **D**: firing rate of inhibitory population, **G**: mean adaptation current, **C**: variance of excitatory firing rate, **F**: variance of inhibitory firing rate, **H**: covariance of the firing rate of the inhibitory and excitatory population). The green curve style represents the stability of the fixed-point (continuous line for stable fix points and dashed lines for unstable fix points). The curve style represents the stability of the fixed-point (continuous line for stable fix points and dashed lines for unstable fix points). The graphics **B** and **E** enlarge the bottom part of their left graphic (A and D). The green crosses are the estimation of the fixed-point using network simulation. The bifurcation points are indicated with a red cross and a letter that means the type of bifurcation (H: Hopf bifurcation and LP: limit point or saddle-node bifurcation).

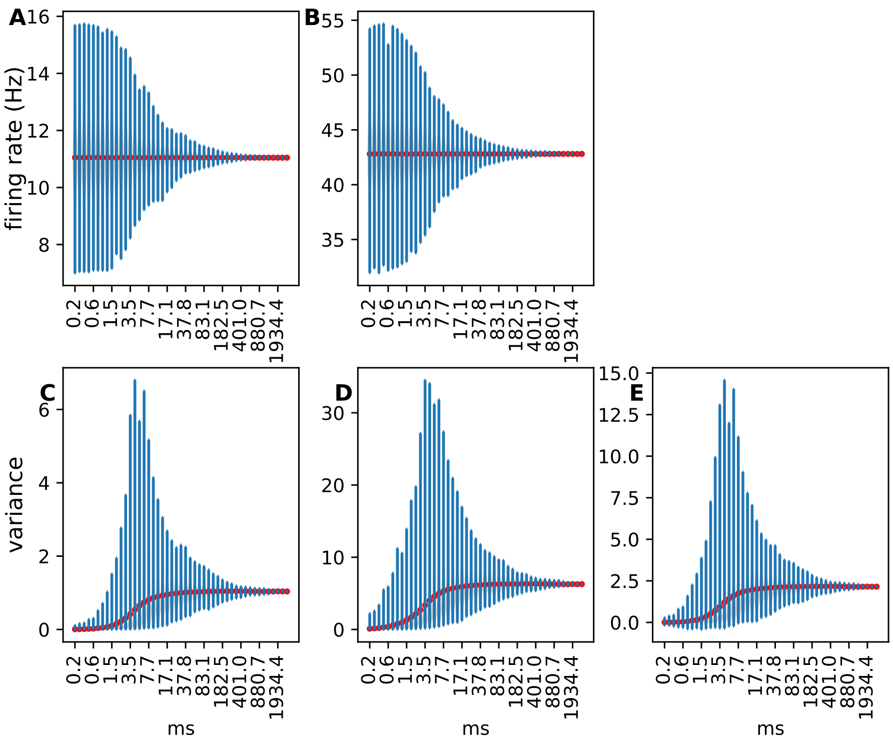


Figure 5: **Variability of the measure of the firing rate** Each violin plot represents the distribution in function of the time windows for the measures associated with the variable of the mean-field, i.e. the mean and variance of firing rate for excitatory and inhibitory. The distribution represents the values of 5000 measures taken randomly over 30 seconds of smoothing instantaneous firing rate with sliding windows of 5 ms. The instantaneous firing rate is recorded from a unique spiking neural network with no adaptation and an excitatory external firing rate of 10 Hz. **A**, **B**: The top graphics represent the mean firing rate of each population. **C**, **D**, **E**: the bottom graphics represent the variance and covariance of these two populations. The left graphics (A and C) are associated with the excitatory population, and the central graphics (C and D) are associated with the inhibitory population. On each graphic, the red line represents the mean of each distribution, and the blue shape represents a rotated kernel density. We can see an exponential decrease in the variability. At the difference of the mean firing rate, the mean of the distribution of the variance and covariance change with the size of the window, and the distribution is not symmetric. The width of the distribution of the variance and covariance increase for a window size of 1.5 ms and 20 ms.

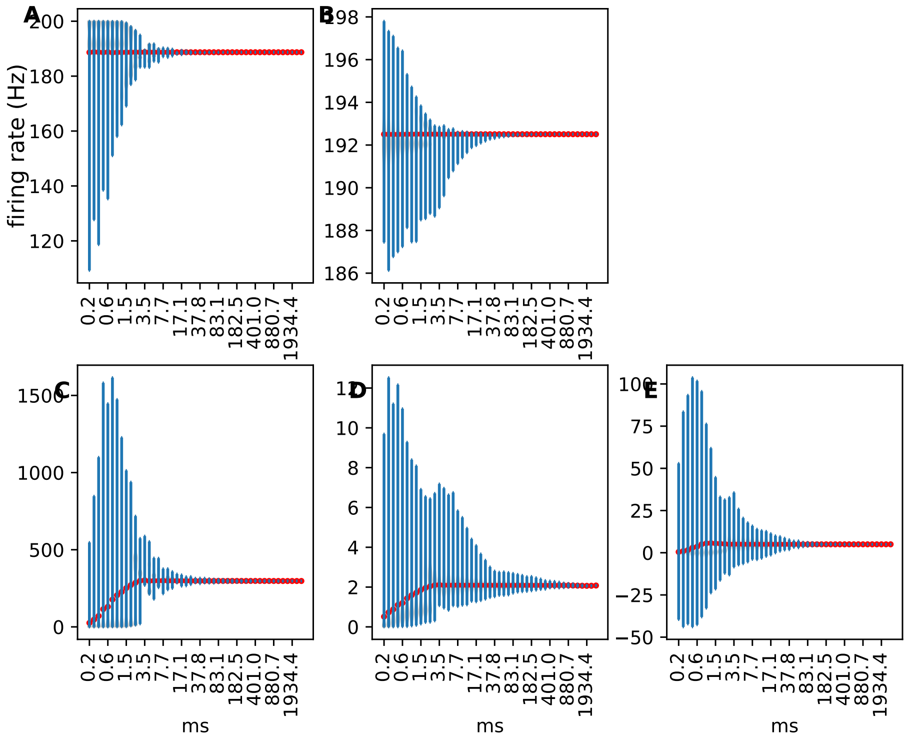


Figure 6: **Variability of the measure of the firing rate** Each violin plot represents the distribution in function of the time windows for the measures associated with the variable of the mean-field, i.e. the mean and variance of firing rate for excitatory and inhibitory. The distribution represents the values of 5000 measures taken randomly over 30 seconds of smoothing instantaneous firing rate with sliding windows of 5 ms. The instantaneous firing rate is recorded from a unique spiking neural network with no adaptation and an excitatory external firing rate of 60 Hz. **A**, **B**: The top graphics represent the mean firing rate of each population. **C**, **D**, **E**: the bottom graphics represent the variance and covariance of these two populations. The left graphics (A and C) are associated with the excitatory population, and the central graphics (C and D) are associated with the inhibitory population. On each graphic, the red line represents the mean of each distribution, and the blue shape represents a rotated kernel density. We can see the variance become smaller for the window's size smaller than the previous figure except for the covariance. At the difference of the mean firing rate, the mean of the distribution of the variance and covariance change with the size of the window. The large variance for small variance is due to the oscillation of the network at a frequency of around 200 Hz.

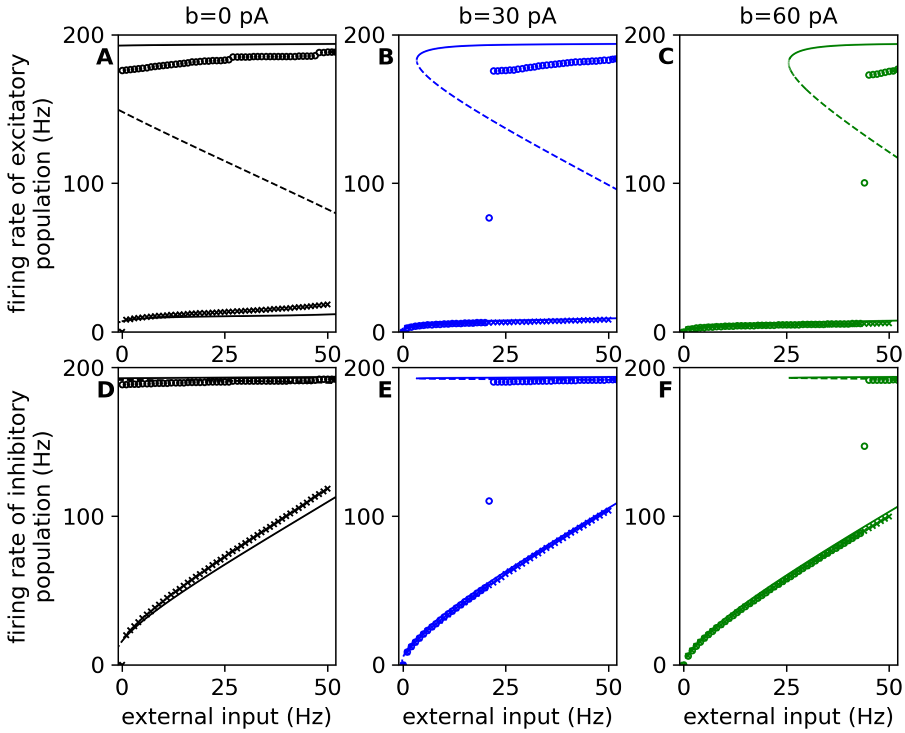


Figure 7: **Estimation and comparison of the high fixed-point A**, **B**, **C**: The top graphics represent the mean firing rate of the excitatory population depending on the external excitatory input value. **D**, **E**, **F**: The bottom graphics represent the mean firing rate of the inhibitory population depending on the external excitatory input. The left graphics (A, D) are for neurons without adaptation, the central graphics (B, E) are for the excitatory population with a spike trigger at 30 pA, and the right graphics (C, F) are for the excitatory population with a spike trigger at 60 pA. The curve and the crosses are the same as the figure 3 (see its caption for more details). The circles represent the mean firing rate over 10 seconds every 10 seconds for one network simulation. During these network simulations, the external excitatory input is reduced to 1 Hz every 10 seconds, from 51 Hz to 0 Hz.

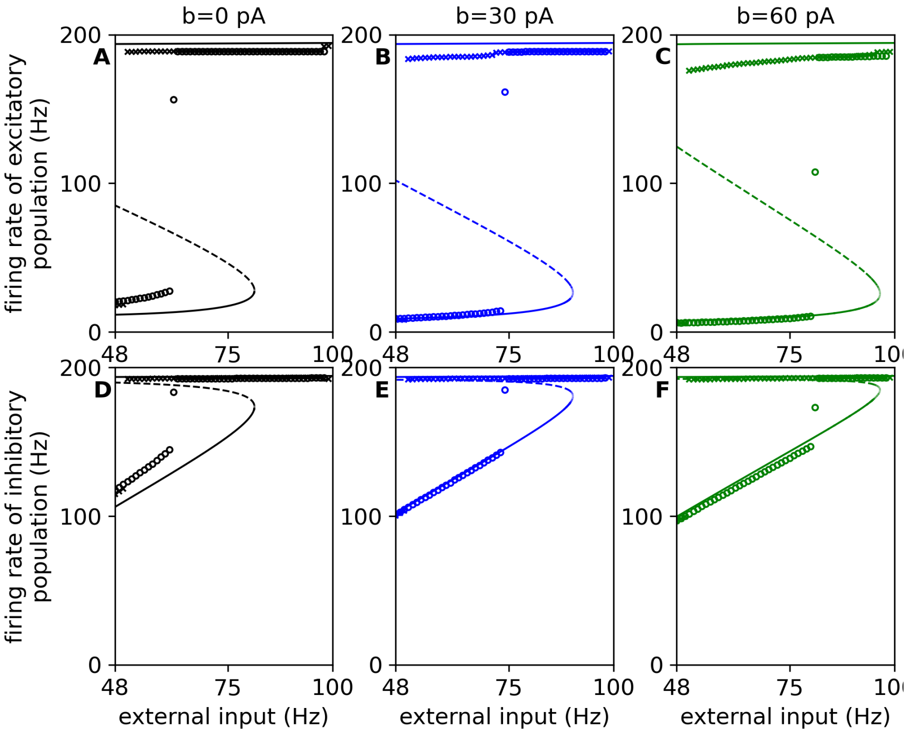


Figure 8: **Estimation and comparison of the low fixed-point A**, **B**, **C**: The top graphics represent the mean firing rate of the excitatory population depending on the external excitatory input value. **D**, **E**, **F**: The bottom graphics represent the mean firing rate of the inhibitory population depending on the external excitatory input. The left graphics (A, D) are for neurons without adaptation, the central graphics (B, E) are for the excitatory population with a spike trigger at 30 pA, and the right graphics (C, F) are for the excitatory population with a spike trigger at 60 pA. The curve and the crosses are the same as the figure 3 (see its caption for more details). The circles represent the mean firing rate over 10 seconds every 10 seconds for one network simulation. During these network simulations, the external excitatory input is reduced to 1 Hz every 10 seconds, from 48 Hz to 99 Hz.

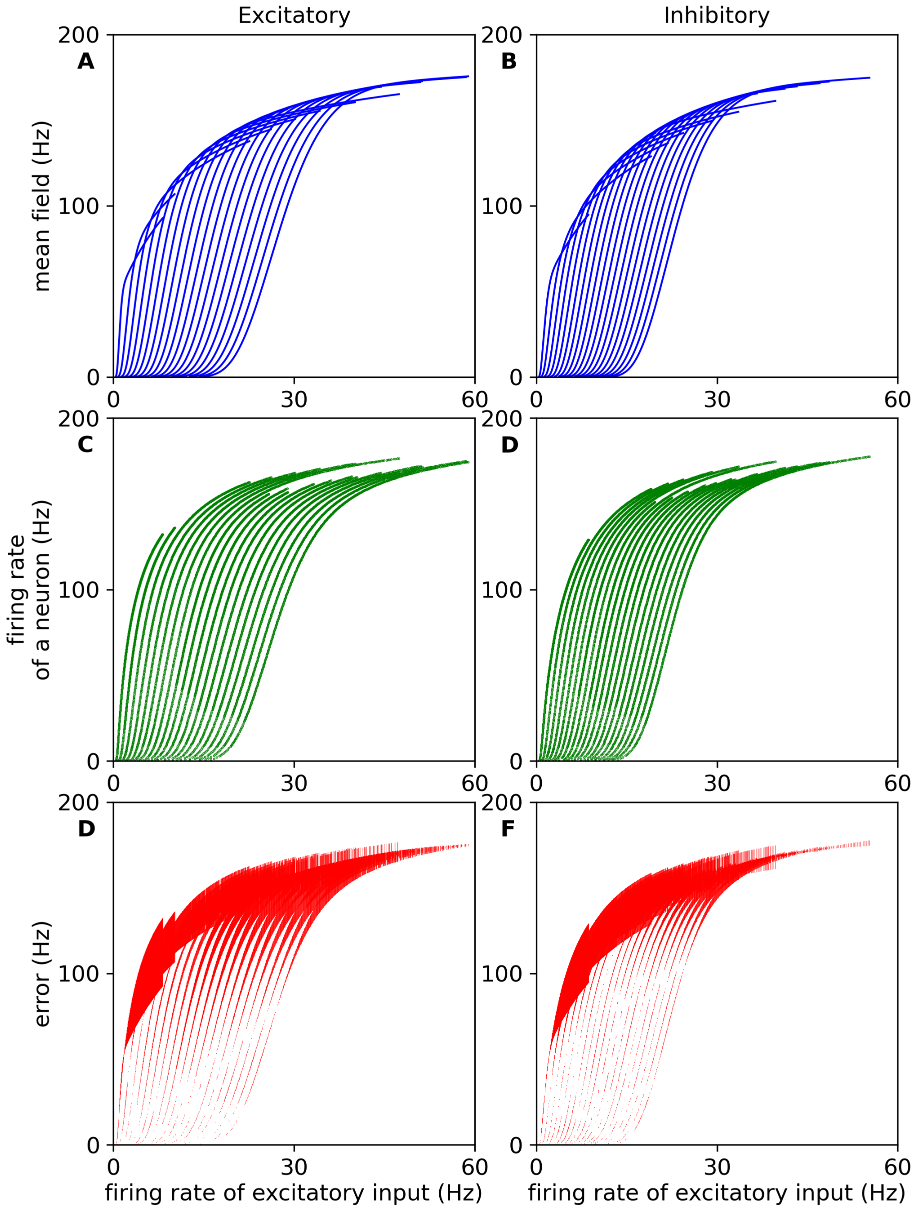


Figure 9: **Fitting of Transfer functions full** This figure enlarges the figure 4. **A**, **C**, **E**: The left part is dedicated to the excitatory neuron. **B**, **D**, **F**, The right part is dedicated to the inhibitory neuron. The top graph (A and B) represents the transfer function of the mean-field fitted to the data. The middle graph (C and D) shows the data used for the fitting, which is the mean firing rate of 50 independent neurons. The bottom graph (E and F) displays the error between the transfer function and the data. Each line is associated with a different value of inhibitory firing rate (20 values evenly distributed between 0 Hz and 40 Hz).

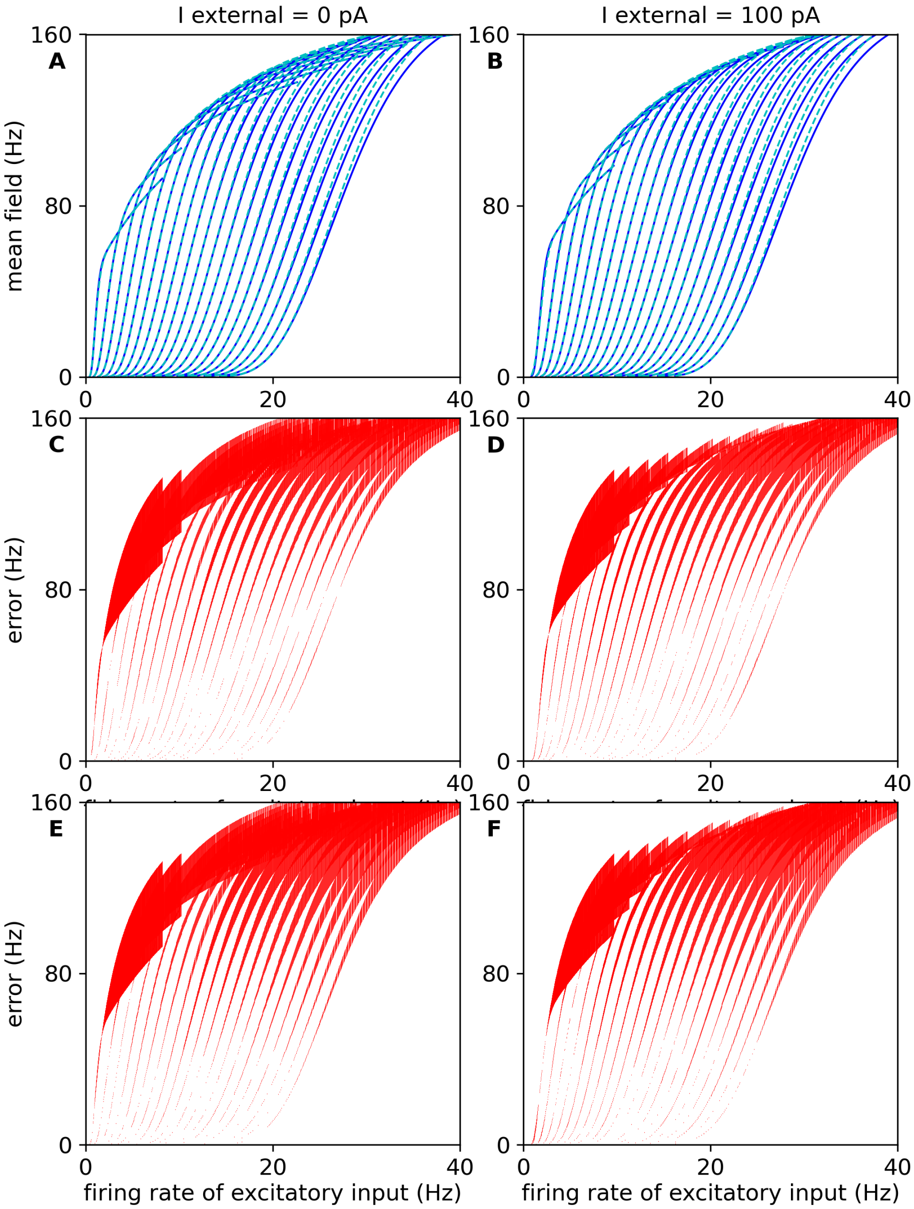


Figure 10: **Estimation of error of the transfer function for neurons in the presence of negative current A**, **B**: The top graphics represent the transfer function fit with (continuous dark blue line) and without (light dashed blue line) data of firing rate with a negative current. **C**, **D**: The middle graphics display the difference between the transfer function fitted with data of negative current. **E**, **F**: The bottom graphics display the difference between the transfer function fitted without the data of negative current. The left graphics is data of the mean firing rate

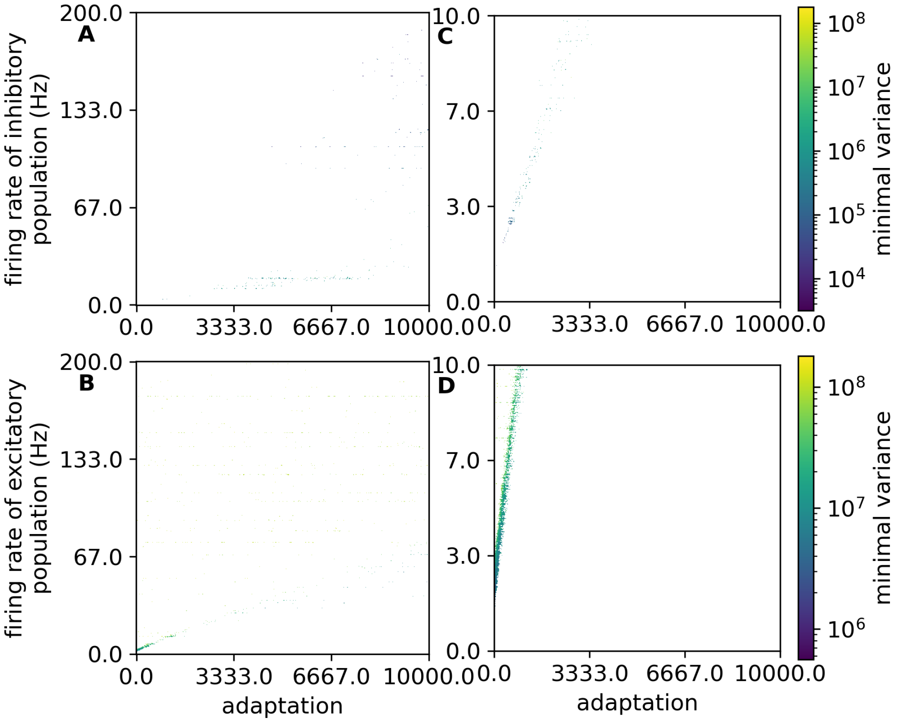


Figure 11: **Negative derivative of the transfer function for zeros firing rate** The graphics display the value of the variance of population A for which the mean firing rate of population B becomes a negative value for a specific value of the mean firing rate of population B and mean adaptive current. **A**, **C**: The top graphic illustrates the variance of the inhibitory population for getting a negative mean excitatory firing rate. **B**, **D**: The bottom graphic illustrates the variance of the excitatory population for getting a negative mean inhibitory firing rate. The right graphics (A and B) enlarge the left graphics (C and D).

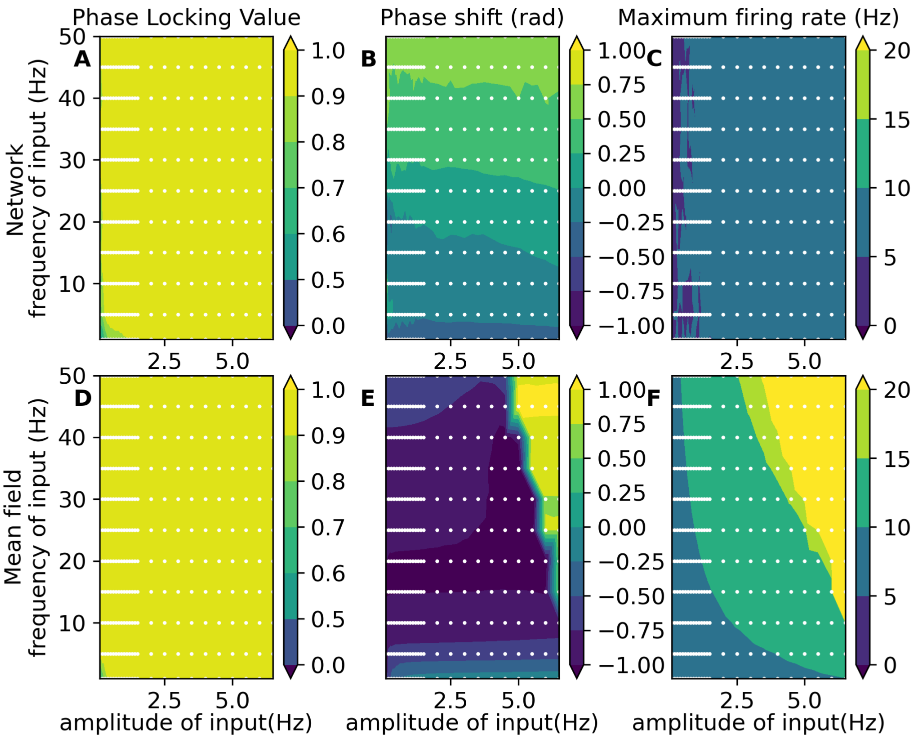


Figure 12: **Comparison between the mean-field and the spiking network in the presence of oscillatory input with a mean rate of 7Hz and adaptation** ($b = 60.0pA$) **A**,**B**, **C**: The top graphics represent the measure of the excitatory population in the spiking neural network. **D**, **E**, **F**: The bottom graphics illustrate the measure of the excitatory mean firing rate of the mean-field. The three measures displayed here are the phase locking values in rad (A and D), the phase shift (B and E) between the oscillatory input and the mean firing rate in rad, and the excitatory population's maximum firing rate in Hz (C and F).

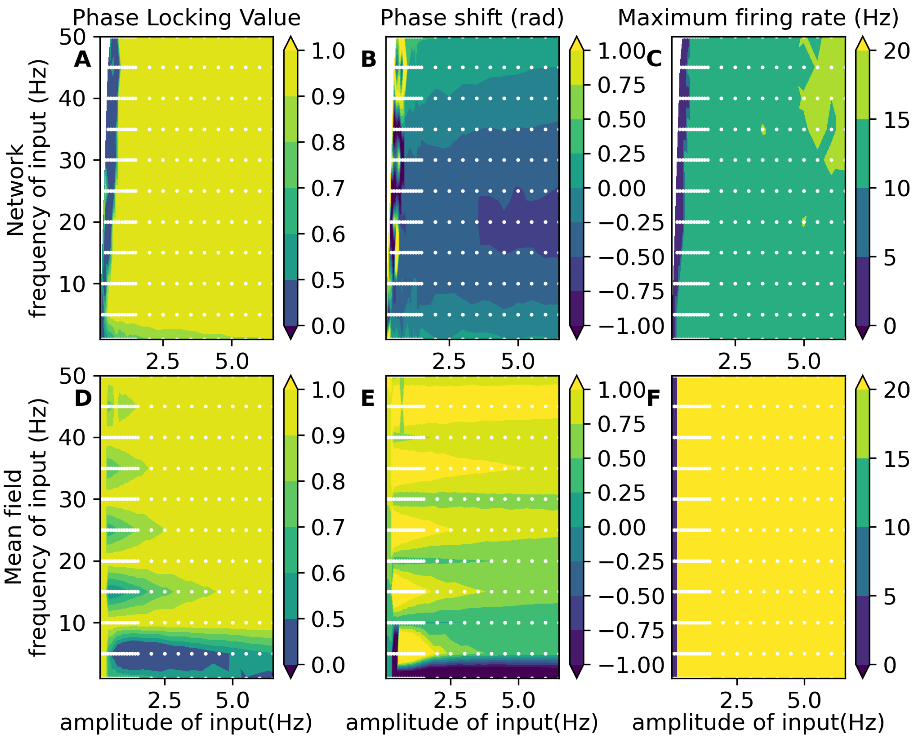


Figure 13: **Comparison between the mean-field and the spiking network in the presence of oscillatory input with a mean rate of 0Hz A**,**B**, **C**: The top graphics represent the measure of the excitatory population in the spiking neural network. **D**, **E**, **F**: The bottom graphics illustrate the measure of the excitatory mean firing rate of the mean-field. The three measures displayed here are the phase locking values in rad (A and D), the phase shift (B and E) between the oscillatory input and the mean firing rate in rad, and the excitatory population's maximum firing rate in Hz (C and F).

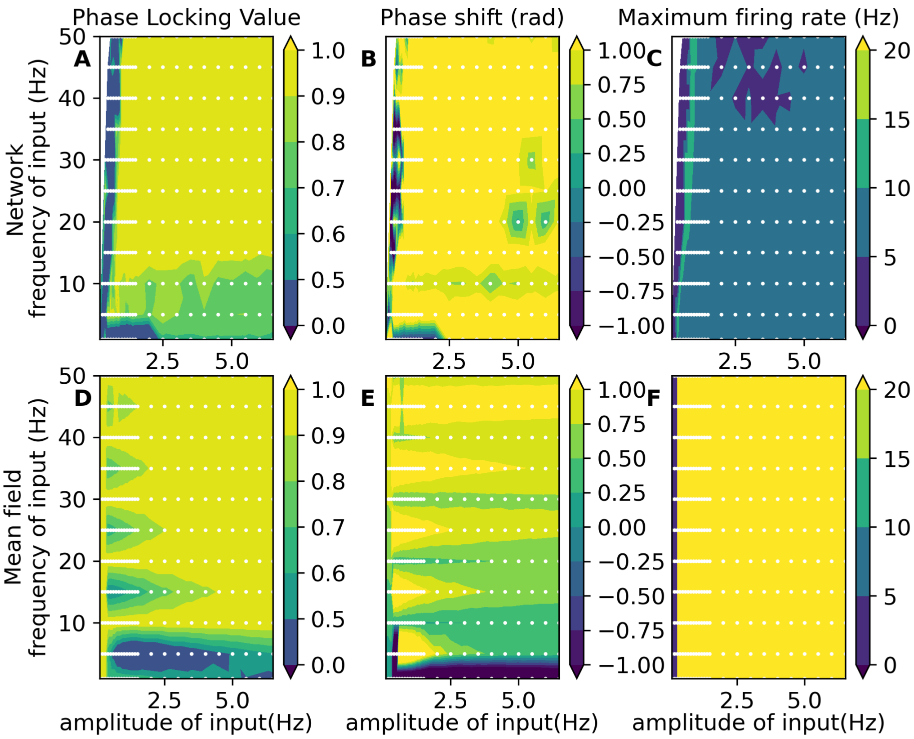


Figure 14: **Comparison between the mean-field and the spiking network in the presence of oscillatory input with a mean rate of 0Hz and adaptation** (**$b = 60.0pA$**) **A**,**B**, **C**: The top graphics represent the measure of the excitatory population in the spiking neural network. **D**, **E**, **F**: The bottom graphics illustrate the measure of the excitatory mean firing rate of the mean-field. The three measures displayed here are the phase locking values in rad (A and D), the phase shift (B and E) between the oscillatory input and the mean firing rate in rad, and the excitatory population's maximum firing rate in Hz (C and F).

[10] Jan Hahne, Sandra Diaz, Alexander Patronis, Wolfram Schenck, Alexander Peyser, Steffen Graber, Sebastian Spreizer, Stine Brekke Vennemo, Tammo Ippen, Håkon Mørk, Jakob Jordan, Johanna Senk, Sara Konradi, Philipp Weidel, Tanguy Fardet, David Dahmen, Dennis Terhorst, Jonas Stapmanns, Guido Trensch, Alexander van Meegen, Jari Pronold, Jochen Martin Eppler, Charl Linssen, Abigail Morrison, Ankur Sinha, Jessica Mitchell, Susanne Kunkel, Rajalekshmi Deepu, Espen Hagen, Tom Vierjahn, Nilton Liuji Kamiji, Robin de Schepper, Pedro Machado, Jasper Albers, Wouter Klijn, Alex Myczko, William Mayner, Pooja Nagendra Babu, Hanjia Jiang, Sebastian Billaudelle, Benedikt S. Vogler, Guilherme Miotto,

Lionel Kusch, Alberto Antonietti, Aitor Morales-Gregorio, Joris Dolderer, Younes Bouhadjar, and Hans Ekkehard Plesser. NEST 3.0, 2021.
